# Nanopore-based sequencing of active DNA replication reveals key principles of metazoan replication fork progression, origin and termination sites

**DOI:** 10.1101/2025.09.26.678856

**Authors:** Dongsheng Han, Cole Shepherd, Mary Lauren Benton, Jared T. Nordman

## Abstract

Balancing replication fork progression and origin usage is essential to maintain genome stability, but measuring replication fork progression rates and origin usage throughout the genome has been challenging. Here, we use nanopore sequencing combined with DNAscent to measure replication fork progression together with origin and termination site usage with single-molecule precision throughout the Drosophila genome with nearly full genome coverage. We find that replication fork progression rates are not uniform throughout the genome. Rather, fork progression is slowest in euchromatin, and this is not correlated with active transcription. Replication origins are also influenced by chromatin, but the exact position of initiation is highly variable and are often several kilobases away from ORC binding sites. Termination sites lack any chromatin or sequence motifs and appear nearly random throughout the genome. By measuring DNA replication dynamics at near full genome coverage, our work reveals key principles of metazoan replication dynamics.

## INTRODUCTION

Complete and faithful replication of the genome is essential to maintain genome stability. DNA replication begins when the Origin Recognition Complex (ORC) binds to thousands of sites throughout the genome and loads a double hexamer of the MCM2-7 replicative helicase complex in a head-to-head orientation^1,2^. Cell cycle dependent kinase activation of the MCM2-7 helicase results in a pair of bidirectional replication forks. Activation of the MCM2-7 helicase and the first synthesis of DNA defines a replication origin. Understanding the factors that control where replication origins are specified and how origin usage is regulated throughout the genome remains an exciting challenge in eukaryotic DNA replication^3^.

While ORC binding in metazoans is largely sequence independent, it is highly influenced by chromatin^4–6^. ORC preferentially binds to nucleosome free regions and directly contributes to the generation of nucleosome free regions^7–11^. ORC binding is not uniform throughout the genome. Rather, there is a higher density of ORC binding sites in early replicating regions of the genome relative to late replicating regions^4,5^. Several chromatin factors are also enriched in ORC binding sites^4,7–11^. While chromatin clearly influences ORC binding, the relationship between metazoan ORC binding sites and origins of replication (start sites of DNA synthesis) is not completely clear. For example, a recent integrative analysis of ORC-binding sites and replication origins in human cultured cells revealed replication origins are often many kilobases from ORC-binding sites^12^.

Multiple population-based approaches have been used to define replication origins on a genome-wide scale^13–19^. While powerful, many of the current techniques for mapping replication initiation sites vary in resolution and concordance. Not only is there variability in origin mapping between techniques, but origin mapping by a single technique, short nascent sequencing or SNS-seq, revealed extensive variation in data sets between laboratories^12^. To circumvent problems with population-level mapping of replication origins and the biases that may be introduced during isolation of nascent DNA, nanopore-based sequencing approaches have been developed to map replication origins in budding yeast and human cells^20,21^. Nanopore-based sequencing of origins has two major advantages to previous origin mapping studies. First, origins can be directly identified from patterns of EdU and BrdU incorporation, alleviating the need for experimental manipulation of samples prior to sequencing. Second, nanopore-based sequencing of origins provides a single molecule view of replication initiation. A recent nanopore-based origin mapping study in human cells has revealed that most origins are dispersed throughout the genome with no defining chromatin or sequence features^21^. While powerful, nanopore-based identification of origins with full coverage of the human genome is cost prohibitive.

Nanopore-based sequencing approaches also have the power to integrate single-molecule measurements of replication fork progression with DNA sequence information. Currently, DNA combing is widely used to measure replication fork progression rates and inter-origin distances ^22,23^. While powerful, DNA combing has notable limitations. First, combing does not provide genomic coordinates for replication tracks, making it impossible to relate fork dynamics to genome-wide chromatin or sequence features^23,24^. Second, DNA combing is a low-throughput and labor-intensive and is prone to technical variability due to inconsistent fiber stretching and analog detection^23,25^. Third, the efficiency of CldU and IdU incorporation can vary between species and cell types, and detection by immunofluorescence can limit both sensitivity and resolution^23,25^. For instance, in Drosophila cultured cells, only CldU is reliably incorporated under standard conditions, limiting the ability to monitor bidirectional fork movement^26^.

Nanopore-based single-molecule sequencing strategies have emerged as all-in-one assays to measure replication fork progression together with initiation and termination sites in a single assay^20,27–31^. Most current studies utilizing nanopore-based sequencing of replication dynamics have been conducted in human or yeast systems^20,27,28,30,32^. Human studies are limited by low genome coverage (typically <10%)^32^,while yeast systems, despite their tractability, lack the complex chromatin environment characteristic of metazoans^27,30,33^. Here we use *Drosophila melanogaster* as model to study metazoan fork progression, origin and termination sites throughout a metazoan genome with near full genome coverage. We find that fork progression rates are slower in euchromatic and early replicating regions compared to heterochromatic and late replicating regions. This difference in fork rate does not correlate with active transcription. We identified 1,513 replication origins and 1,984 termination sites. Replication origins are influenced by chromatin but the exact position where initiation occurs is variable while termination sites have no defining chromatin or sequence-related elements.

## RESULTS

### Nanopore can measure replication initiation, termination, and fork progression in Drosophila cultured cells

To measure DNA replication dynamics throughout the entire Drosophila genome with nearly full genome coverage and DNA sequence resolution, we adapted and optimized an approach that integrates Oxford Nanopore long-read sequencing with DNA thymidine analog labeling of nascent DNA^20,27,28,30,32^. Specific for Drosophila S2 cells, we refined the timing and concentration of nucleotide analog pulses and optimized DNA extraction protocols to maximize yield and DNA integrity. Cells were sequentially labeled with 5-ethynyl-2’-deoxyuridine (EdU) for 7 minutes, followed by a 15-minute chase with 5-bromo-2’-deoxyuridine (BrdU) (Fig. 1A). Both analogs can be distinguished from thymidine during Nanopore sequencing based on their unique current signal profiles^20,27,28,30,32^. After extracting genomic DNA, we prepared PCR-free libraries and performed sequencing using the Oxford Nanopore platform (Fig. 1A). We then used the machine learning-based software DNAscent^34^ to accurately identify EdU, BrdU, and thymidine incorporation patterns along individual DNA molecules. In wild-type (WT) cells, both EdU and BrdU were efficiently incorporated into nascent DNA (Fig. 1B). We were able to capture various aspects of replication dynamics—including fork directionality, initiation, and termination events— at single-molecule resolution (Fig. 1B).

**Fig. 1.**
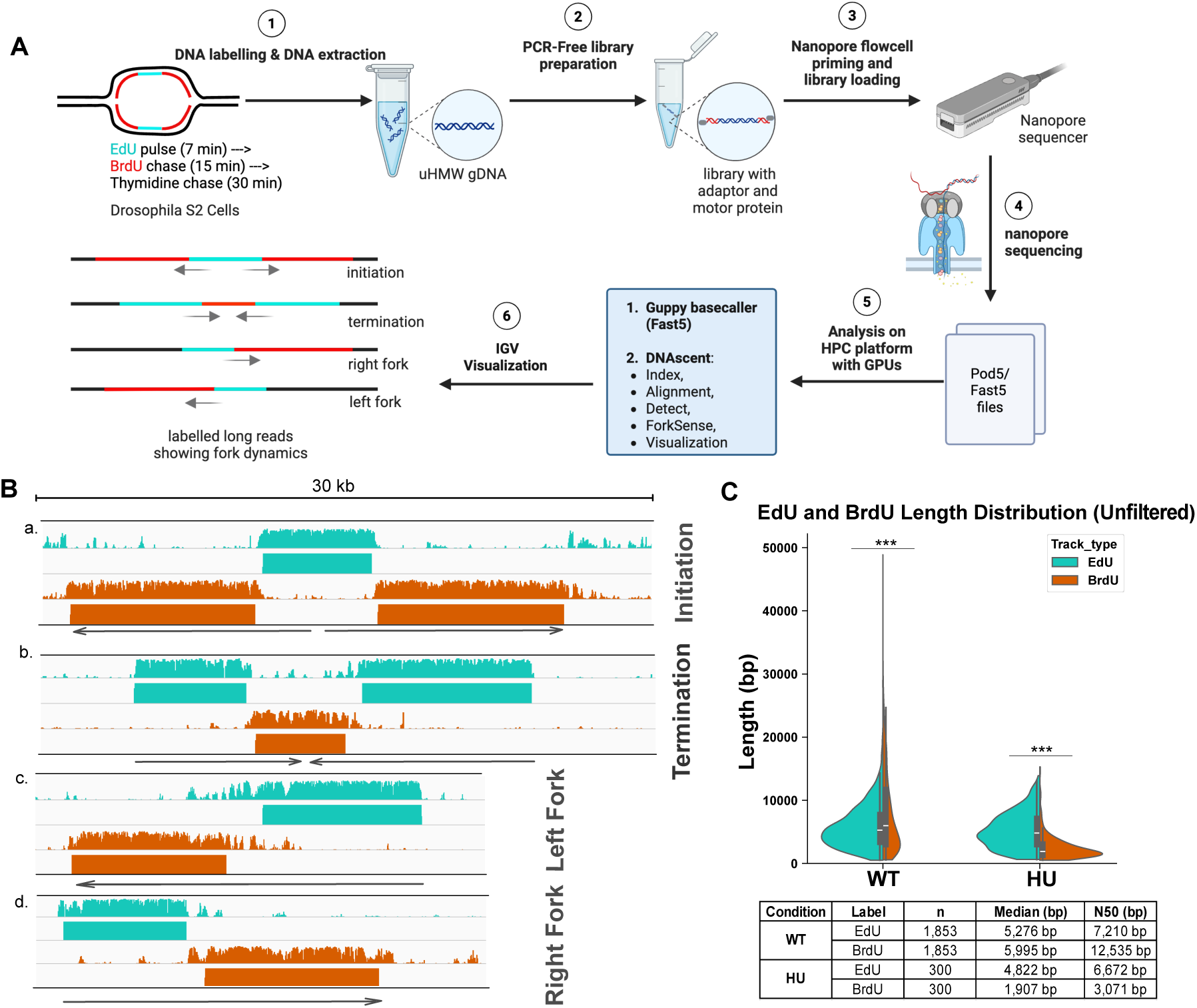
Nanopore-based replication fork capture and analysis. **(A)** Schematic overview of the workflow for nanopore-based sequencing of replication dynamics in Drosophila S2 cells. S2 cells are subjected to sequential pulses of EdU (cyan) and BrdU (red) to label nascent DNA. After DNA extraction, ultra-high molecular weight genomic DNA (uHMW gDNA) is prepared for nanopore sequencing without PCR amplification. The prepared library is loaded onto the Nanopore flowcell and sequenced. The raw data is then analyzed on a high-performance computing (HPC) platform Guppy basecaller and the DNAscent software suite for signal processing and visualization. The final output consists of labelled long reads that show the dynamics of replication forks. **(B)** Examples of labelled long reads showing different replication fork dynamics. Cyan tracks correspond to EdU-labelled DNA, and orange tracis correspond to BrdU-labelled DNA. The grey line with arrows indicates the direction of the replication forks. The scale bar represents 30 kilobases (kb). **a.** Example of a replication initiation event. **b.** Example of a replication termination event. **c.** Example of a leftward moving fork. **d.** Example of a rightward moving fork. **(C)** Unfiltered violin plots showing the length distribution of EdU- and BrdU-labelled segments in untreated and hydroxyurea-treated (HU) cells. The horizontal lines within the violins indicate the median. The table below the plot shows the number of reads (n), the median length, and the N50 length for each condition. Statistical significance was calculated using a two-sided Mann–Whitney U test. ***p<0.001.

To validate this method, we exposed cells to the DNA replication inhibitor hydroxyurea (HU) at the start of the BrdU chase^35^. This experimental design allowed EdU tracks to reflect fork progression before replication stress, while BrdU tracks captured fork behavior under stress. In untreated cells, BrdU-labeled tracks were significantly longer than EdU-labeled tracks, with median lengths of 5,995 bp (N50: 12,535 bp) and 5,276 bp (N50: 7,210 bp), respectively. Upon HU treatment, BrdU track lengths were dramatically reduced (median: 1,907 bp; N50: 3,071 bp), while EdU-labeled tracks remained relatively unchanged (median: 4,822 bp; N50: 6,672 bp) (Fig. 1C). These results demonstrate that Nanopore-based approach is highly sensitive to replication stress and enables high-resolution, single-molecule analysis of replication fork dynamics in Drosophila cultured cells.

### Nanopore-based sequencing can measure replication dynamics with near full genome coverage in Drosophila

Having established nanopore sequencing to measure replication dynamics in Drosophila cultured cells, we wanted to extend this approach to measure replication dynamics with higher genome coverage. To achieve this, we applied the same labeling and extraction approach but sequenced the resulting library across two PromethION flow cells. This approach yielded a total of 25,057 unfiltered replication forks, 1,513 replication origins and 1,984 replication termination sites. To ensure data quality, we excluded replication fork reads when the BrdU tracks extending to the terminal ends of the molecule^29,34^. After filtering, we retained 11,449 high-quality labeled molecules, which we consider high-confidence replication forks. EdU-labeled segments of these high confidence forks had a mean length of 6,223bp, a median of 5,512bp and an N50 of 7,668 bp (Fig. 2A). In contrast, BrdU-labeled segments were much longer, with a mean of 10,525bp, a median of 8,815bp and an N50 of 14,543bp. Based on the N50 length over a 15-minute labeling period, this translates to a fork progression rate of 969.5bp/min in S2 cells. The ratio of BrdU to EdU segment lengths had a median of 1.50 and a mean of 2.13 (Fig. 2B), consistent with the relative durations of our EdU and BrdU pulses.

**Fig. 2.**
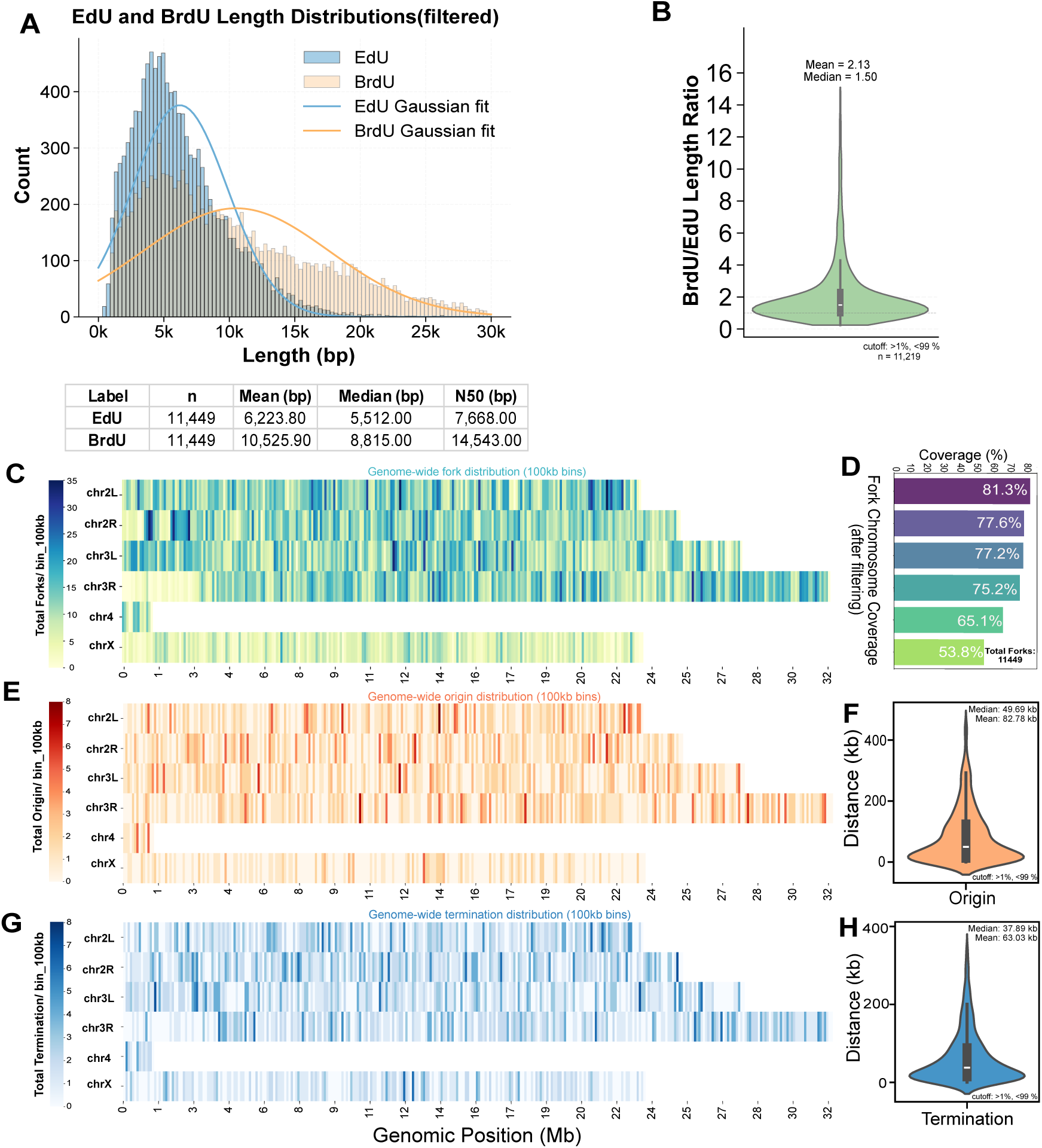
Nanopore based sequencing of replication dynamics achieves near genome-wide coverage. **(A)** Filtered length distributions of EdU- and BrdU-labeled segments. The histogram shows the count of segments at various lengths in base pairs (bp). Gaussian curves (orange and blue lines) are fitted to the BrdU and EdU distributions, respectively. The table below provides the number of reads (n), mean length, median length, and N50 length for each labeled population. **(B)** Violin plot showing the distribution of the BrdU-to-EdU length ratio for 11,219 replication tracks. The plot indicates the relative length of the BrdU-labeled track compared to the EdU-labeled track for per single labeled DNA molecule. The mean and median values for the ratio are shown above the plot. The plot excludes values outside the 1st and 99th percentiles. **(C)** Genome-wide distribution of replication forks. The heatmap shows the density of replication forks across different chromosomes (*chr2L, chr2R, chr3L, chr3R, chr4, chrX*) in 100kb bins. The color intensity represents the fork density, with lighter colors indicating lower density and darker colors indicating higher density. The percentage of chromosomal coverage is shown in the table to the right. **(D)** Table summarizing the chromosomal coverage (%) for the genome-wide fork distribution shown in panel C. **(E)** Genome-wide distribution of replication origins. This heatmap shows the density of replication origins across the chromosomes in 100kb bins. The color intensity represents the origin density. **(F)** Violin plot showing the inter-origin distances. The mean and median values are indicated above the plot. The plot excludes values outside the 1st and 99th percentiles. **(G)** Genome-wide distribution of replication terminations. This heatmap shows the density of replication termination events across the chromosomes in 100kb bins. The color intensity represents the termination density. The plot excludes values outside the 1st and 99th percentiles. **(H)** Violin plot showing the inter-termination distances. The mean and median values are indicated above the plot. The plot excludes values outside the 1st and 99th percentiles.

Next, we sought to determine whether the number of replication forks, origin and termination sites we obtained was sufficient to achieve full genome coverage, and if so, whether that coverage was uniformly distributed across the genome. We found nearly complete coverage of the major chromosome arms (Fig. 2C). For example, prior to filtering, *chr2L* exhibited 94.3% coverage (23.5 Mb), while *chrX* showed slightly lower coverage at 78.4% (23.6 Mb) (Fig. 2D). After filtering, fork coverage across chromosomes ranged from 81.3% (*chr2L*) to 53.8% (*chrX*) (Fig. 2D). When visualized in 100kb bins, fork distribution was non-uniform across the genome (Fig. 2C), with regions of high and low fork density. The density of replication forks was strongly correlated with DNA copy number differences in Drosophila S2 cells^36^. We next identified replication origins and termination sites based on the spatial arrangement of EdU and BrdU labels within individual molecules (Fig. 1A). To ensure uniformity, we took the midpoint of each origin and termination site and added 100bp to each side. We identified a total of 1,513 origin sites and 1,984 termination events throughout the genome. Replication origins were unevenly distributed across the genome (Fig. 2E), with a median inter-origin distance of 49.69kb and a mean of 82.78kb (Fig. 2F), consistent with previously reported inter-origin distances in *Drosophila* cultured cells using DNA fibers^37^. Termination sites also exhibited variable genomic density (Fig. 2G), with a median inter-termination distance of 37.89kb and a mean of 63.03kb (Fig. 2H). Similar to replication forks, replication origins and termination sites were not uniformly distributed across the genome. Instead, both features exhibited regions of local enrichment and depletion (Fig. 2E and 2G), reflecting the heterogeneous nature of replication initiation and completion across different chromosomal contexts and chromosomal copy number changes in Drosophila S2 cells^36,38^. Taken together, data from only two PromethION flow cells can provide enough labeled molecules to achieve near-complete coverage of the Drosophila genome, enabling robust genome-wide profiling of replication initiation, progression, and termination events.

### Replication fork progression is influenced by chromatin

Like replication tracks measured by DNA combing, DNAscent revealed substantial variation in BrdU track lengths, potentially reflecting differences in replication fork speeds across the genome. To assess whether variation in track length is associate with chromatin state, we mapped BrdU tracks on the established nine-state Drosophila chromatin model^38^, including updated regions that are predominantly repetitive and heterochromatic (State 0) (Supplemental Fig. 1A). During the conversion of the nine-state chromatin model from the dm3 to the dm6 genome assembly, regions newly present in dm6 but absent in dm3 were assigned to the heterochromatin state, based on these regions being predominantly composed of mappable repetitive sequences and are typically annotated as heterochromatin in Drosophila^39^. Analysis of BrdU track lengths across chromatin states revealed significant differences (Fig. 3A). Unexpectedly, chromatin states associated with heterochromatin or repressive features— including State 0 (Dm3-not-covered), State 6 (Polycomb-repressed), and State 7 (Heterochromatin)—exhibited longer BrdU tracks than euchromatin-associated states (States 1– 5)^38^. State 9, which lacks defined histone modifications^38^, also showed longer BrdU tracks relative to most euchromatic states (Supplemental Fig. 1A). To more easily compare fork progression between broad chromatin compartments, we grouped the chromatin states into euchromatin (States 1–5) and heterochromatin (States 0, 6, 7, 8). Consistent with individual state analysis, BrdU track length was significant longer in heterochromatin (median= 63,558 bp) than in euchromatin (median= 56,258 bp). To account for potential bias driven by the DNA molecule size during filtering (Supplemental Fig. 1B, C), we normalized the BrdU track length to total read length for each read. Normalized BrdU track lengths revealed the same trend with longer BrdU tracks in heterochromatic relative to euchromatic states (Supplemental Fig. 1D).

**Fig. 3.**
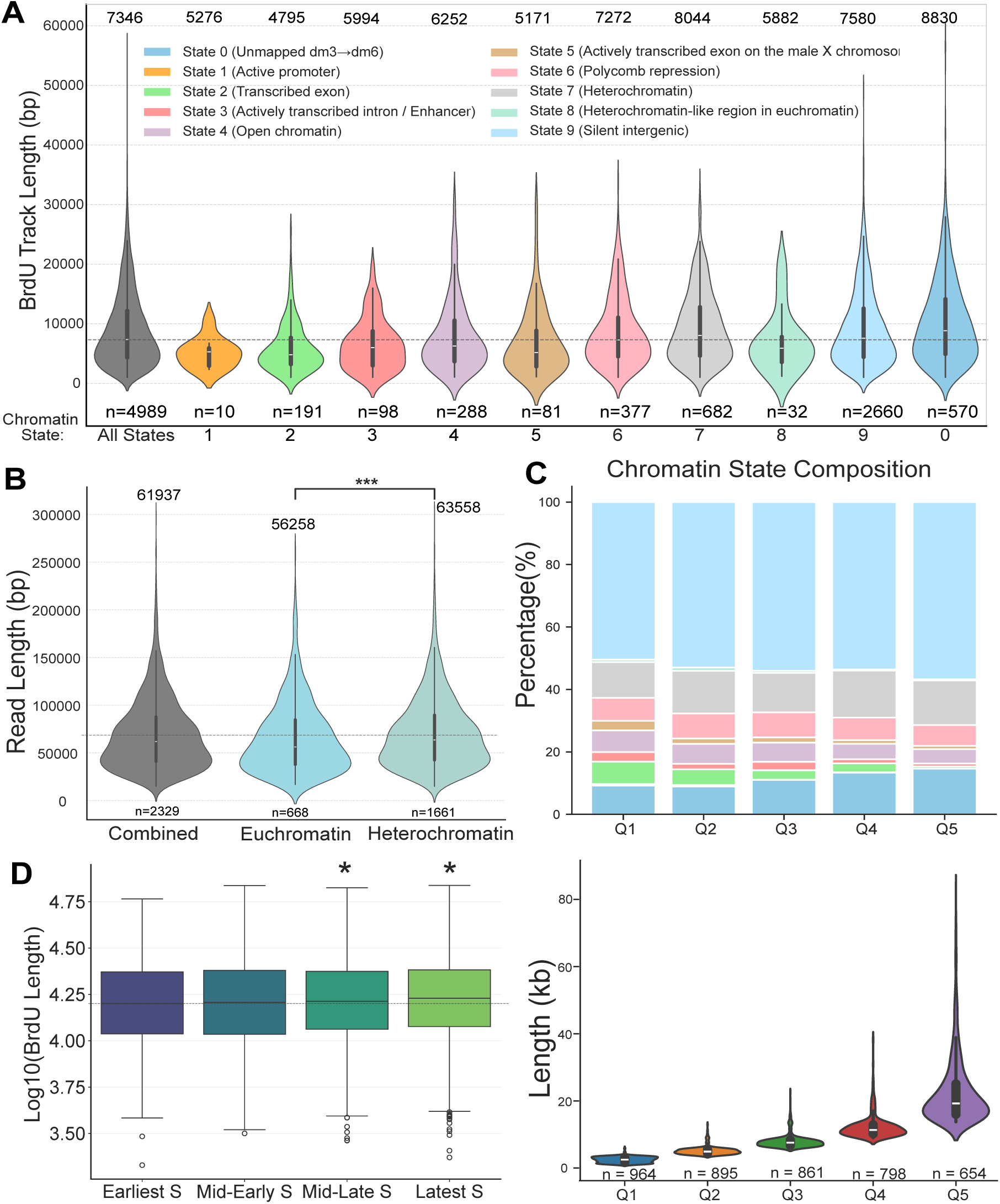
Distribution of BrdU track lengths across chromatin states and S-phase stages. **(A)** Violin plot showing the distribution of BrdU track lengths (in base pairs, bp) across ten different chromatin states. The first violin represents all reads (“All States”), while the remaining violins correspond to individual states defined in Supplemental Fig. 1A. Heterochromatin-associated states (0, 6, 7, and 9) exhibit longer BrdU tracks than euchromatin-associated states (1–5). The number of tracks (n) for each state is shown below the plot. The median BrdU track length (bp) for each is above the plot. **(B)** Violin plot comparing the distribution of BrdU read lengths (bp) between broad chromatin domains. BrdU tracks are grouped into euchromatin (States 1–5), heterochromatin (States 0, 6, 7, 8), and an all-reads “combined” group. The number of reads (n) for each group is shown below the plot. Statistical significance was determined using a two-sided Mann-Whitney U test. ***p<0.001. **(C)** (Upper) Stacked bar chart depicting the percentage composition of chromatin states within BrdU track length quartiles (Q1–Q5). The y-axis represents the proportion of each quartile comprised of different chromatin states, color-coded as in panel A. (Bottom) Violin plot showing the distribution of BrdU read lengths (kilobases, kb) within each quartile. The number of reads (n) per quartile is indicated below the plot. **(D)** Box plot showing the distribution of log_10_-transformed BrdU read lengths across distinct S-phase stages (Earliest S, Mid-Early S, Mid-Late S, Latest S). Boxes represent the interquartile range (IQR), with the median indicated by the central line. Outliers are plotted as individual points. Statistical comparisons between each stage and the earliest S phase were performed using a two-sided Mann-Whitney U test. *p < 0.05.

To independently validate this observation, we grouped BrdU tracks into quintiles by length (Fig. 3C, bottom panel) and analyzed the chromatin state composition in each quintile. The longest track quintiles (Q4 and Q5) were enriched for heterochromatin-associated states, such as State 0 and State 7. In contrast, euchromatic states such as State 2 and State 5 were overrepresented in the shortest quintiles (Q1 and Q2). When normalized to the genomic representation of each state, this trend persisted and heterochromatin states were consistently overrepresented among the longest BrdU track quintiles, while euchromatin states were enriched among the shortest (Supplemental Fig. 1E). Interestingly, State 9 was also overrepresented in longer track quintiles (Fig. 3C, S3E), suggesting that defined heterochromatin chromatin marks are not necessarily the driver of increased fork progression rates.

Given that replication timing (RT) is tightly linked to chromatin state, we next asked whether fork progression differs between early- and late-replicating regions^6,40–43^. We first compared BrdU lengths across four discrete RT groups using existing RT data sets generated in S2 cells (Fig. 3D) ^6^. The BrdU track lengths in “Mid-Late S” and “Latest S” regions were significantly longer than those in earlier replicating regions (Fig. 3D). To assess this relationship more rigorously, we assigned a RT score to each BrdU-labeled track and measured its correlation with track length. This analysis revealed a weak but statistically significant negative correlation (Pearson’s R = –0.04; p = 1.8 × 10⁻⁶), indicating that replication fork progression is slower in early-replicating regions. We confirmed that, as expected, early-replicating regions were predominantly euchromatic, while late-replicating regions were enriched for heterochromatin (Supplemental Fig. 1G). Taken together, these data indicate that replication fork progression is faster in later-replicating, heterochromatin-rich regions when compared to early replicating euchromatin. Importantly, this trend has recently been observed in human cells suggesting a more universal principle in control of replication fork progression rates^32,44^.

### Differences in fork progression across chromatin states are largely independent of transcription

Given the significant differences in replication fork speed observed across distinct chromatin states, we investigated whether these differences could be attributed to transcriptional activity. Since replication-transcription conflicts shape fork speed and stability^45,46^, we hypothesized that slower fork speeds observed in euchromatic regions might be related to higher levels of gene transcription. To test this, we examined the relationship between BrdU track length (as a proxy for fork speed) and active transcription using published PRO-seq data in Drosophila S2 cells^47^. First, we compared activated transcription levels across BrdU track length quintiles. As shown in Fig. 4A, mean RPKM values did not differ significantly between the shortest (Q1) and longest (Q5) BrdU track groups. A cumulative distribution analysis of transcription levels between all quintiles revealed substantial overlap, consistent with minimal differences between BrdU track length and active transcription (Fig. 4B). We next performed a linear correlation analysis between BrdU track length and transcription levels, which revealed no correlation between BrdU track length and transcription levels (Pearson’s R = −0.01, p = 2.91 × 10^⁻1^) (Fig. 4C). To determine whether chromatin context might influence this relationship, we performed correlation analyses within specific chromatin subtypes. We restricted our analysis to BrdU tracks located entirely within either euchromatin (States 1–5) or heterochromatin (States 0, 6, 7, 8)^38^. A slight correlation between transcription levels and BrdU track length was observed in euchromatin (R = −0.07, p = 3.28 x 10^-2^, n = 817) but no significant correlation was determined in heterochromatin (R = −0.02, p = 6.11 x 10^-1^, n = 762) (Fig. 4C).

**Fig. 4.**
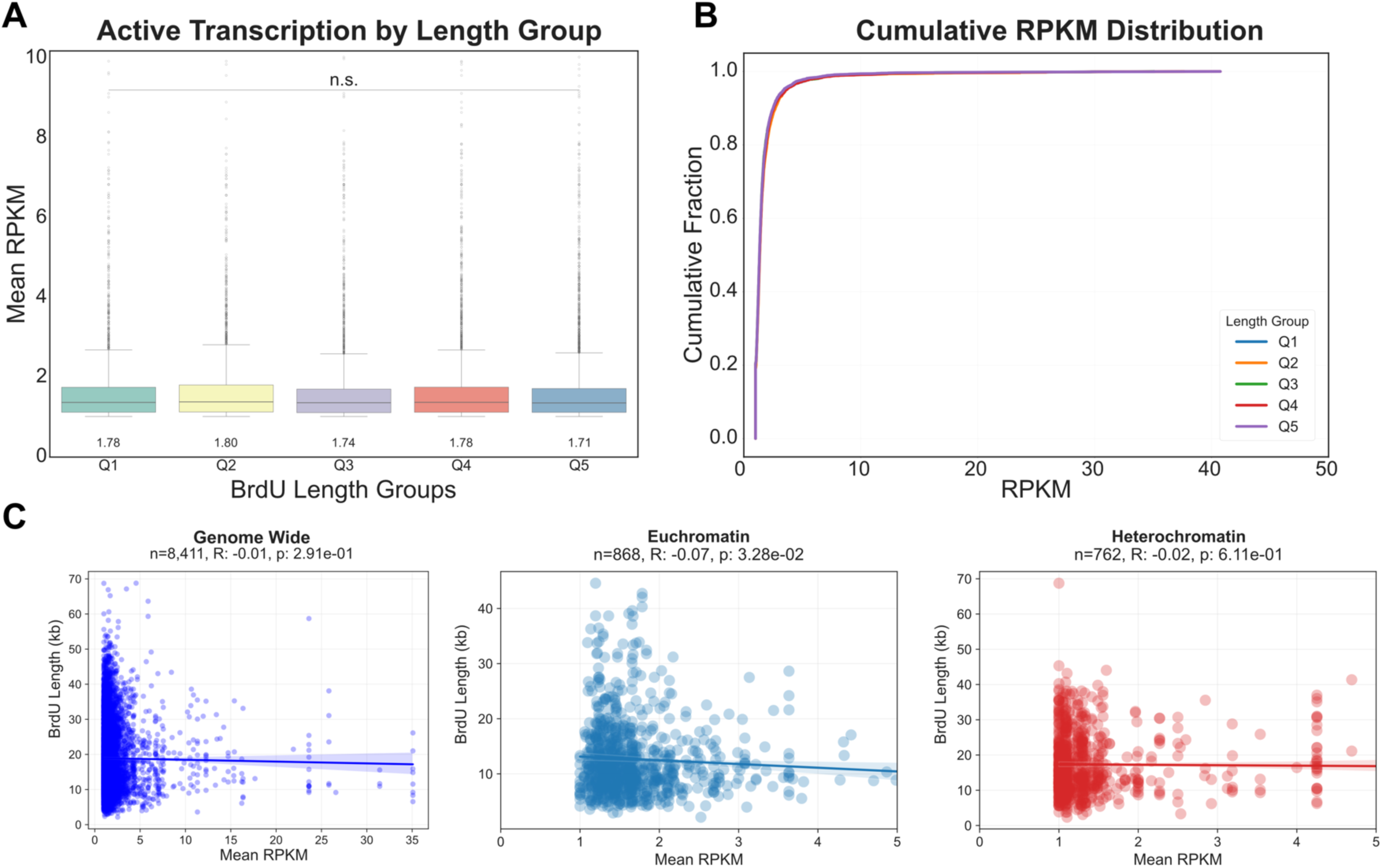
Differences in fork progression rates across chromatin states do not correlate with transcript levels. **(A)** Box plot showing the distribution of mean transcription activity (in reads per kilobase million) across five BrdU track length quintiles (Q1-Q5). Mean RPKM values (listed under the box plots) did not differ significantly between the shortest (Q1) and longest (Q5) track groups (n.s. – not significant). Statistical differences between groups were tested using Mann–Whitney U tests. **(B)** Cumulative distribution function (CDF) plot showing the cumulative fraction of BrdU tracks as a function of transcription activity for each length quintile (Q1-Q5). The substantial overlap of the curves for all five length groups indicates that overall active transcriptional profiles are similar regardless of BrdU track length. **(C)** Scatter plots showing the correlation between BrdU track lengths and active transcription. The analyses are shown for all nonzero genes in three different genomic contexts.

We asked whether the orientation of transcription relative to replication fork direction might influence fork progression. For regions where transcription and replication were oriented counter directional (head-on), we observed no significant correlation between BrdU track length and head-on transcription (Pearson’s R = −0.006; p = 7.58 x 10^-1^; n = 2,425). Similarly, no significant correlation was observed in regions with co-directional transcription and replication (Pearson’s R = −0.032; p =8.7 x 10^-2^; n = 2,799) (Supplemental Fig. 2A-B).

Taken together, these results demonstrate that transcription is unlikely to explain the observed differences in replication fork progression rates across chromatin states. Thus, additional features associated with these chromatin environments are likely to drive differential fork progression rates.

### Nanopore sequencing revealed non-uniform replication fork directionality genome-wide

PCR-free Nanopore-based sequencing preserves strand information, enabling analysis of replication fork directionality. In Drosophila germ line stem cells, histone H3 is asymmetrically distributed to daughter cells through an unknown mechanism^48,49^. One possibility is that large scale unidirectional replication of the genome drives the asymmetric distribution of histones in these germ line stem cells^50^. To test if DNAscent can be used to measure biases in fork directionally throughout the genome, we measured the replication fork directionality throughout the genome in 100kb window sizes (Fig. 5A). We found the vast majority of windows were replicated with both leftward and rightward forks (Fig. 5A-B). While we did observe a small percentage of windows that did show bias in fork directionality (Supplemental Figure 3C), there was no large-scale unidirectional replication of the genome in S2 cells (Fig. 5A). Nanopore-based sequencing strategies should also be able to capture replication dynamics of the mitochondrial genome. The mitochondrial genome is replicated through a strand-displacement mechanism and leftward and rightward forks are expected to map to a specific template strand (Fig. 5C). Therefore, we mapped replication tracks to the mitochondrial genome in a strand-specific manner. We detected strand-specific patterns consistent with the strand-displacement model of mtDNA replication^51^(Fig. 5D). The small size of the mitochondrial genome and the different rate for fork progression in the mitochondrial genome, will require separate optimization for ideal measurements of fork progression rates. Regardless, Nanopore-based sequencing can be used to measure both nuclear and mitochondrial DNA replication dynamics.

**Fig. 5.**
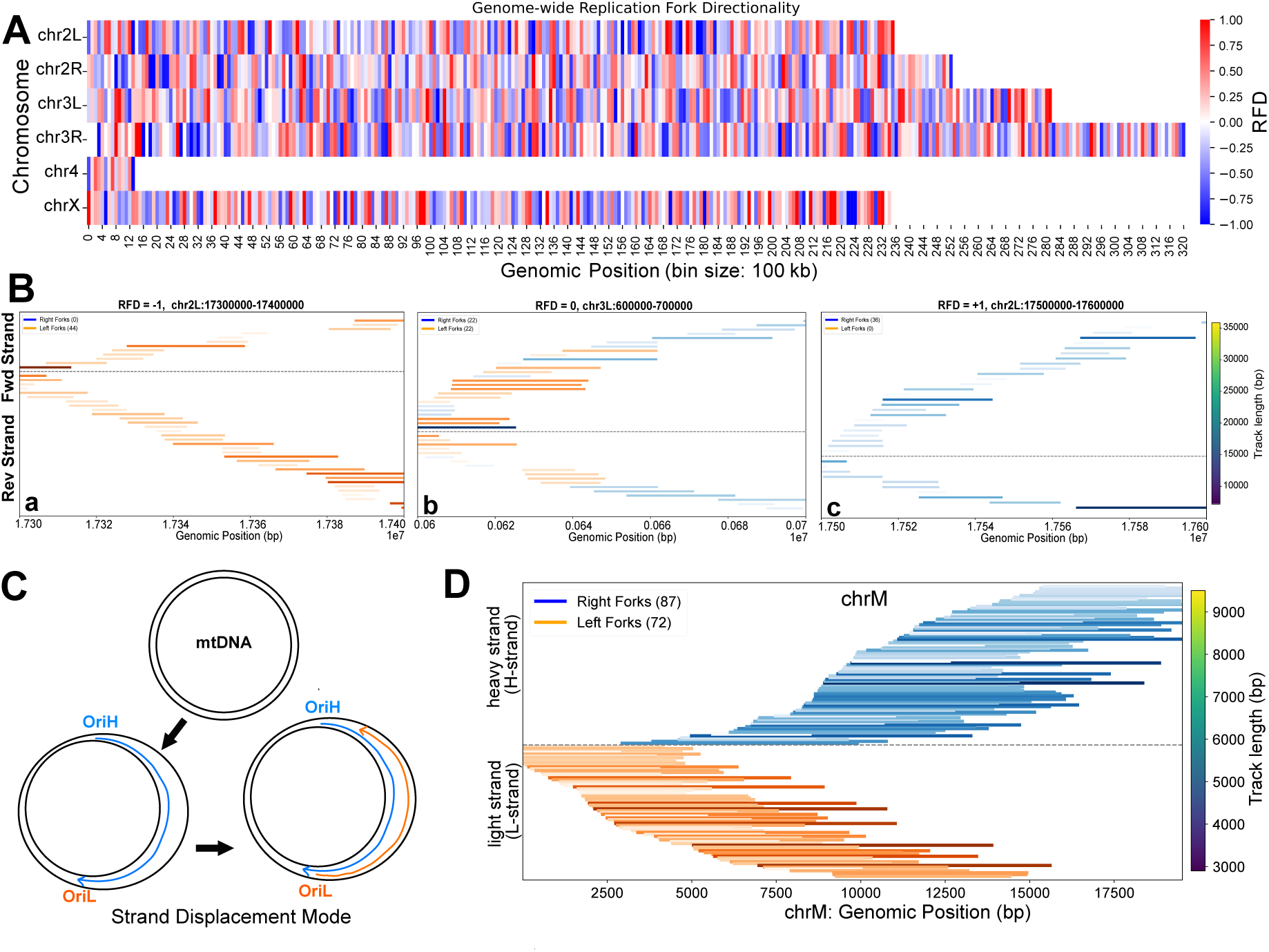
Genome-wide and mitochondrial DNA replication fork directionality. **(A)** Heatmap showing the genome-wide distribution of Replication Fork Directionality (RFD) in 100 kilobase (kb) bins across Drosophila chromosomes. The color scale (red to blue) represents the RFD value, with positive values indicating a bias towards one direction and negative values indicating a bias towards the other. The lack of large red or blue blocks indicates no substantial unidirectional replication across the genome. **(B)** Screenshots of individual 100kb window showing a. leftward fork bias, b. leftward and rightward fork replication, c. rightward fork bias. **(C)** Quantification of fork bias in 100kb windows throughout the genome. RFD of 1 = 100% rightward forks and RFD of −1 = 100% leftward forks. **(D)** A schematic of the mitochondrial DNA replication through the strand displacement mechanism. **(E)** Visualization of strand-specific replication patterns on mitochondrial DNA (mtDNA). Reads are plotted on either the forward (fwd) or reverse (rev) strand, with each bar representing a single read. The length of each bar corresponds to its genomic position on the chromosome (*chrM*), and its color indicates the BrdU length (in base pairs, bp) as shown by the color bar.

### Origins are influenced by chromatin and are often distant from ORC sites

DNAscent identified 1,513 replication origins genome-wide, with a median inter-origin distance of 49.69 kb (Fig. 2F). This distance is close to a reported inter-origin distance of ∼40kb measured in Drosophila cultured cell by DNA fiber analysis^37^. Therefore, we were able to analyze origin usage throughout the entire Drosophila genome at near full coverage. To assess whether origin density is influenced by replication timing, we compared origin density between early and late replicating regions. Replication origins are initially licensed through ORC-dependent MCM loading^52–54^ and the density of ORC2 binding sites higher in early replicating regions relative to late replicating regions (Supplemental Fig. 4B)^4,6^. Similar to ORC binding sites, origin density was also higher in early domains than in late domains (1.45 vs 0.73 origins per 100 kb; Fig. 6A), though the difference was smaller than that observed for ORC2 binding (Supplemental Fig. 4B). Next, we examined whether chromatin state influences origin usage. Like the densities in early and late replicating regions, the density of origins was higher in euchromatic regions of the genome, which was also true for ORC binding sites (Fig. 6B; Supplemental Fig. 4C). Together, these results indicate that origin density is influenced by replication timing and/or chromatin structure.

**Fig. 6.**
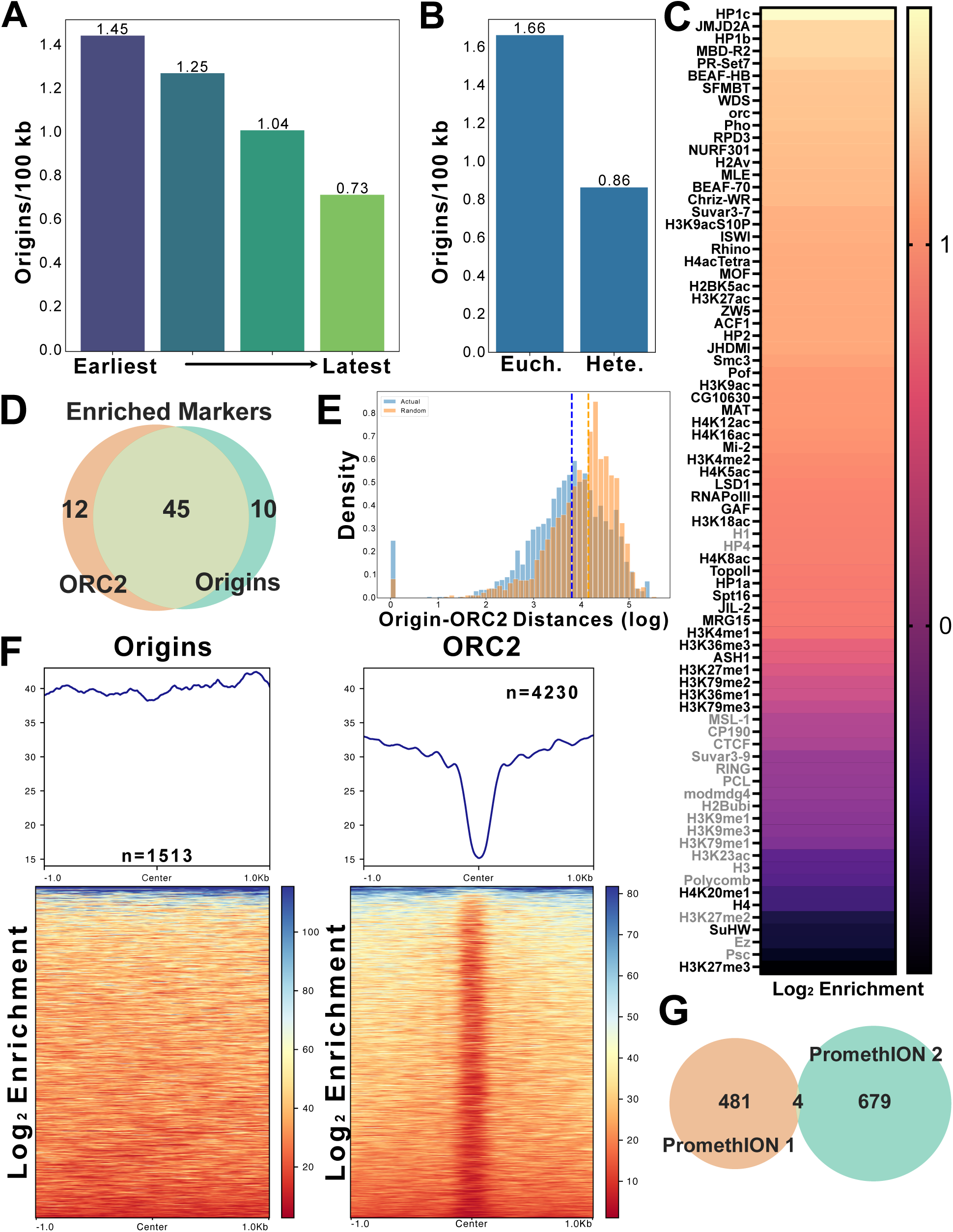
Replication origin distribution, chromatin context, and usage at the single-molecule level. **(A)** Bar chart showing replication origin density across early and late replicating domains. **(B)** Bar chart comparing replication origin density between euchromatin and heterochromatin. **(C)** Heatmap showing the log_2_ enrichment values of chromatin factors and histone modifications at replication origins. Log_2_ enrichment values were calculated based on the observed overlap relative to the expected overlap for each chromatin factor. Black indicates statistically significant enriched or depleted factors (FDR <0.05%). Grey indicates non-significant enriched or depleted factors. **(D)** Venn diagram illustrating the overlap between chromatin factors significantly enriched at replication origins and those significantly enriched at ORC2 binding sites. **(E)** Histogram comparing the distribution of distances between replication origins and the nearest ORC2 peak (“Actual”) or a randomized control (“Random”). **(F)** Heatmap visualizing the chromatin accessibility by MNase-seq at replication origins and ORC2 sites. **(G)** Venn diagram showing the overlap of identified replication origins (size=200bp) between two biological replicates (“PromethION 1” and “PromethION 2”).

The differential origin density between euchromatin and heterochromatin suggests that chromatin structure could modulate origin usage. Since ORC binding is influenced by chromatin features such as nucleosome positioning and chromatin-associated factors^55–58^, we asked whether specific chromatin factors are positively or negatively associated with replication origins. To this end, we measured the frequency of overlap between origins and chromatin-associated factors with ChIP data available through modENCODE^38,59^. To determine if enrichments were more or less than expected by chance, we shuffled regions throughout the genome and calculated the amount of overlap for each permutation^60,61^. Many chromatin factors showed significant enrichment at origins (Fig. 6E). Interestingly, the majority of these chromatin factors are also significantly enriched at ORC2 binding sites^7^ (Fig. 6F). Importantly, more ORC2 binding sites overlapped with origins than would be expected by chance (Fig. 6F; Supplemental Fig. 4F).

While ORC2-binding sites and origins could share common chromatin features, there were also differences. The median distance between an origin and the nearest ORC2 biding site was 6.309kb, which was closer than distance expected by chance (median – 13.099 kb) (Fig. 6E). This is consistent with a recent meta-analysis of initiation sites and ORC binding sites in mammalian cells^12^ and suggests a functional relationship between ORC2 binding and replication initiation. ORC binding is enriched in nucleosome-depleted regions^6,62^ (Fig. 6F). In contrast, replication origins mapped by single-molecule assays show only a weak correlation with nucleosome-free regions identified by population-based methods (Fig. 6F). This indicates that, unlike ORC sites, initiation sites do not require well-defined nucleosome-free regions.

We next examined sequence features associated with replication origins. We found no specific sequence motifs enriched at origins (Supplemental Fig. 4E), nor any consistent bias in AT or GC content (Supplemental Fig. 4F). Instead, origins preferentially localize to promoters and transcription termination sites (TTS), while being depleted within gene bodies, including exons and introns (Supplemental Fig. 4G). Taken together, this suggests that ORC-loaded MCMs are likely influenced by transcription, resulting in origins many kilobases away from where MCMs were initially loaded^6,21,62–64^.

In two independent experiments, we identified 485 and 683 origins. Surprisingly, only four origins were shared between these replicates (Fig. 6G). Even when the overlap window was expanded tenfold, only 36 origins overlapped between the two biological replicates (Supplemental Fig. 4H). We then pooled all origins from multiple individual replicates (n = 1,513) and found that only 29 origins that appeared more than once. Assuming an inter-origin distance of 40kb^37^, we expect 3,593 origins in the mappable genome (see Methods). Random sampling of 3,593 sites would result in 277 shared sites based on the size of our data set (see Methods). Furthermore, only ∼20% of replication origins were found in clusters (defined as three or more origins within a 40kb window – Supplemental Fig. 4I). Together with our results showing that origins lack any sequence bias or motifs, we conclude that origin usage is influenced by chromatin, but the precise sites of initiation are dispersed throughout the Drosophila genome. Finally, we compared our single-molecule mapping of replication origins with a published population-based SNS-seq origin mapping dataset performed in the same Drosophila cell line^65^. Surprisingly, we found little overlap between the two datasets and fewer overlapping sites than expected by chance (Supplemental Fig. 4D). Thus, the majority of origins we identified by a single molecule assay have been missed by a population-based approach.

### Termination sites do not correlate with chromatin features or sequence motifs

Population-based studies have not been able to identify precise sites of replication termination^13,14^. This is because termination happens in zones that cannot be precisely defined by lagging-strand mapping methods^13,14^. Nanopore has been used to map termination events in budding yeast and human cells, however, genome wide analysis of termination sites has only been performed in budding yeast^20^. Here it was found that termination sites are more dispersed than previously thought based on population-based studies. DNAscent identified 1,984 termination sites that were distributed throughout the entire Drosophila genome with a median inter-termination distance of 37.89 kb (Fig. 2G-H), similar to the median inter-origin distance of 49.69 kb.

In contrast to origins, there was an increased number of termination sites in late replication regions of the genome (Fig. 7A). The distribution of termination events as measured by inter-termination distance, however, was not significantly different between heterochromatin and euchromatin (Fig. 7B). To ask if the density of termination events was influenced by chromatin, we quantified the density of termination sites in euchromatin and heterochromatin found nearly no difference (Fig. 7C). We used a random shuffling method to determine if specific chromatin factors are enriched at termination sites. While there were some enrichment and depletions, none of these reached the level of significance (FDR <0.05; Fig. 7C). Furthermore, no AT/GC sequence bias or sequence motif was found at termination sites (Supplemental Fig. 5A-B). Termination sites between biological replicates showed very little overlap (Supplemental Fig. 5D-E) and only 16 out of 1,984 termination sites were mapped more than once from the combined data sets. Therefore, we conclude that termination sites are potentially random throughout the genome with no defining chromatin or sequence features.

**Fig. 7:**
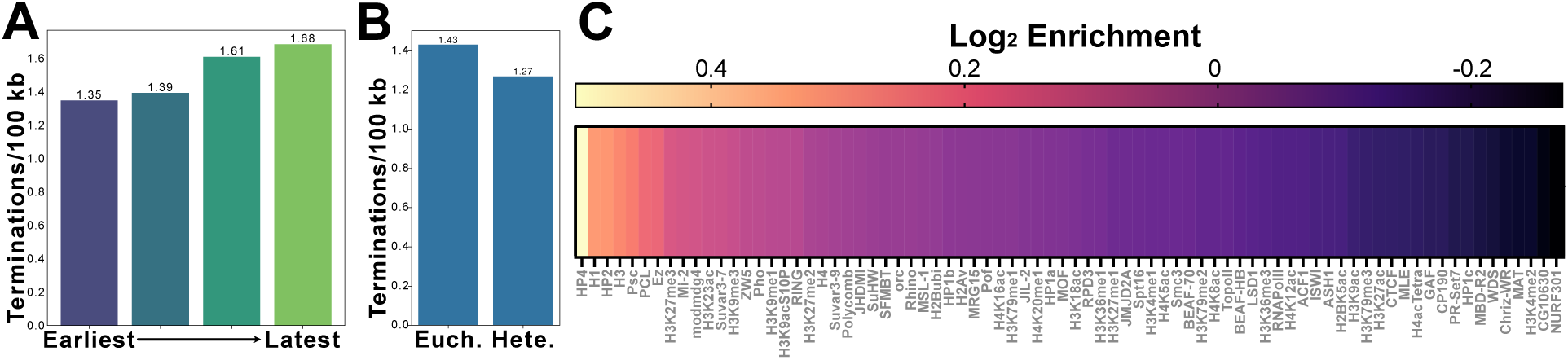
Termination events appear to be stochastic. **(A)** Bar chart comparing the density of termination events (peaks per 100 kb) between early and late replicating regions. **(B)** Bar chart comparing replication origin density between euchromatin and heterochromatin. **(C)** Heatmap showing the log_2_ enrichment values of chromatin factors and histone modifications at replication termination sites. Log_2_ enrichment values were calculated based on the observed overlap relative to the expected overlap for each chromatin factor. Black indicates statistically significant enriched or depleted factors (FDR <0.05%). Grey indicates non-significant enriched or depleted factors.

## DISCUSSION

Nanopore-based sequencing strategies have the potential to answer long-standing questions about metazoan DNA replication that have been difficult to address due to technical limitations^20,21,27,30^. Here, we have used Nanopore-based sequencing of active DNA replication using DNAscent to understand key principles of metazoan replication fork dynamics, origin and termination site selection with single molecule precision^32,34^. Given the genome size of *Drosophila melanogaster*, we were able to define single molecule replication events that cover nearly the entire genome with two PromethION flow cells. Thus, we have provided the first single molecule analysis of replication dynamics covering nearly an entire metazoan genome. DNA combing has been the gold standard method to measure rates of replication fork progression throughout the genome^20,27–31^. While powerful, this method is unable to provide sequence-level information. Therefore, it is impossible to know if track length variability in DNA combining experiments is caused by random fluctuations in nucleotide incorporation rates or, more interestingly, reflect differences in fork progression rates throughout the genome driven by biological constraints to fork progression. We were able to address this issue by combining BrdU and EdU pulses with DNAscent, generating single molecule measurements of fork progression throughout the Drosophila genome. To our surprise, we found that replication fork progression is slower in euchromatic and early replicating regions of the genome relative to heterochromatic and late replication portions of the genome. Nanopore-based measurements of replication fork progression in budding yeast found no difference in fork progression rates throughout the genome^30^. In contrast, our work in Drosophila and recent Nanopore-based measurements of fork progression in human cells, have revealed that fork progression is slower in early replicating regions of the genome relative to late replication regions^32,44^. Furthermore, quantification of replication fork speed in single cells have also found the same trend^44^. It is likely that the higher order chromatin structure found in metazoans, and lacking in budding yeast, could contribute to differences in fork rates throughout the genome. For example, budding yeast lacks large blocks of pericentric heterochromatin and the linker histone H1^66,67^.

The driving force causing reduced fork rate speeds in euchromatin relative to heterochromatin remains unresolved. One potential driver of fork slowing in early and euchromatic regions are replication-transcription conflicts^45,68,69^. In fact, single-cell measurements of fork speeds in human cells revealed that transcription is at least partially responsible for the slowing of fork progression in early replicating regions of the genome^44^. Our work in Drosophila, however, suggests that transcription is unlikely to contribute to fork slowing in euchromatic regions of the genome. We were unable to find a correlation between transcript abundance and fork progression that would suggest transcription shapes the rate of fork progression throughout the genome. Nor did we see a difference in fork progression rates when forks were traversing transcription units in head-on vs. codirectional orientation, as would be predicted if slowing was driven by replication-transcription conflicts^45,68,69^. The difference between our results and single-cell sequencing in human cells could be driven by single molecule vs. population level approaches or differences between human and Drosophila DNA replication. An independent study of fork progression rate in human cells, however, identified the same slowing of fork progression in early replicating regions that appears independent of transcription^29^. If not transcription, what is driving slower replication in early and euchromatic regions of the genome? One possibility is topological stress due to more frequent origin firing in early replicating and euchromatic regions of the genome could slow fork progression. Importantly, we do observe an increased density of origins in these regions.

DNAscent allows for the identification of replication forks together with replication initiation and termination sites ^20,21,27,30^. Thus, we were able to define replication origins from single-molecule replication events throughout the entire genome. Several strategies have been used to define replication origins from using population-based approaches. Nanopore-based sequencing, however, provides a single-molecule view of replication initiation allowing a much more precise view of where initiation actually starts in the genome. By performing this analysis in Drosophila, we were able to provide coverage of nearly the entire genome, thus revealing key features of metazoan replication initiation. Surprisingly, of the >1,500 replication initiation sites we identified, very few were identified more than once. Given the genome coverage, number of events identified and expected inter-origin distance, we would have expected to identify individual origins more than once. This suggests that, at the single molecule level, where initiation starts is relatively stochastic. Importantly, Nanopore-based sequencing of origins in human cells and a meta-analysis of initiation sites in human cells came to a similar conclusion^12,70^.

While DNA sequence does not influence where initiation starts, chromatin context does. For example, we found a higher density of origins in early and euchromatic regions of the genome relative to late and heterochromatic regions. Furthermore, we were able to identify chromatin features that were statistically enriched at origins. Importantly, many of these features were similar to features enriched at ORC binding sites^7^. While there was some overlap between origins and ORC binding sites, the median distance between an origin and ORC-binding site was closer than expected by chance. A recent meta-analysis of ORC-binding sites and origins in human cell lines found the vast majority of origins were >1kb away from an ORC binding site^12^. Thus, physical separation of ORC binding sites and replication origins seems to be a consistent trend in metazoan replication. While there is a formal possibility that specification of initiation sites is independent of ORC, MCM distribution is shaped by transcription and Cyclin E activity. Therefore, we proposed that ORC-loaded MCMs moved by transcription and other chromatin forces are stochastically activated by DDK. Thus, the MCMs are a moving target rather than a static complex waiting to be activated. This would explain the distance between ORC and initiation sites and why actual sites of initiation, defined by a single molecule approach, are dispersed throughout the genome.

We also identified 1,984 termination sites. To our knowledge, there has been no formal analysis of metazoan termination sites at the single molecule level. Population-based approaches such as OK-seq or SNS-seq cannot define precise sites of replication termination because termination occurs in zones between initiation sites^71–73^. Nanopore-based strategies have been used to define termination sites in budding yeast and revealed they were often dispersed between replication origins^27,30^. We failed to identify any sequence bias or chromatin signature associated with termination sites in metazoan. Thus, replication termination events occur stochastically throughout the genome without sequence specificity or a chromatin signature.

Nanopore sequencing technology has the potential to revolutionize our understanding of DNA replication dynamics. This single molecule, long-read sequence technology can reveal properties of fork progression, origin and termination site usage throughout the genome in a single assay. Our work reveals that nanopore-sequencing based approaches can measure replication dynamics with almost full coverage of a metazoan genome. Given the only requirement is the incorporation of BrdU and EdU, this technology can be adapted to many organisms, cell types and tissues. Furthermore, as a telomere-to-telomere genome assembly becomes available for the Drosophila genome, our data sets can be re-analyzed to define replication dynamics in regions of the genome that have been difficult to map, including satellite sequences.

## METHODS

### Cell culture

S2 cell stocks were maintained in T225 Flasks with room temperature Schneider S2 Media supplemented with 10% fetal bovine serum (FBS), 100 U/mL Penicillin, and 100 µg/mL Streptomycin. Cells were passaged 1:5 dilution (1-part old media: 5-part final volume) when flasks reached 90-100% confluency (fully confluent). New stocks were thawed and maintained after 30 passages.

### DNA Labeling and Validation for Nanopore Sequencing

24-hours prior to labeling, a fully confluent flask was passaged 1:2 or 1:3 to achieve 60-80% confluency at the time of labeling to ensure cells were actively replicating. The two flasks of cells resulting from this split were labeled at the same time and combined during a later step. The day of labeling, old media was removed and immediately replaced with 25 mL of new media supplemented with 10µM EdU. Cells were returned to the 25°C incubator for 3 or 7 minutes for labeling. After the appropriate time had passed, media was removed, and the cells were gently washed three times with 10mL of 1x Phosphate Buffer Saline (1x PBS). Cells were then incubated with 25mL of new media supplemented with 10µM BrdU. For HU treatment, 1mM was added to the BrdU media. Cells were returned to the 25 °C incubator for 6 or 15 minutes for labeling. After the appropriate time had passed, media was removed, and the cells were gently wash three times with 10mL of 1x Phosphate Buffer Saline (1x PBS). The cells were then incubated with 25mL of new media supplemented with 50µM thymidine. Cells were returned to the 25 °C incubator for 20 minutes before harvesting. Cells from pairs of flasks labeled at the same time were combined into a single 50mL conical and 200µL of cells were set aside on ice for later counting and labeling validation. The remaining cells were pelleted at 600xg for 5 minutes and the old media was removed. The cells were then washed via resuspending in 10mL 1x PBS and subsequent pelleting at 600xg for 5 minutes. This wash step was repeated once. After washing, the cells were resuspended into 1mL 1x PBS, transferred to a 1.5mL Eppendorf tube, and pelleted at 600xg for 5 minutes. The supernatant was discarded and the cells pellets were stored at −80 °C for up to two weeks until DNA extraction and sequencing.

### Extraction of uHMW gDNA for Nanopore Sequencing

uHMW gDNA extraction and library preparation was conducted as a variation on New England Biolab’s adaption for their T3050L Kit for Oxford Nanopore Sequencing. EB+ was prepared by supplementing 1mL of Monarch Elution Buffer 2 with 2µL of 10% Triton X-100. 100 million previously labeled and frozen cells were resuspended to a concentration of 50 million cells per 100uL 1x PBS and divided into two 1.5mL Eppendorf for parallel extraction. The cells were lysed using 100µL Nuclei Prep Buffer and 50µL of 10mg/mL RNase A and mixed at room temperature for two minutes. Nuclei Lysis Solution was prepared using 200µL Nuclei Lysis Buffer and 80µL 1mg/mL Proteinase K. The nuclei were then lysed using 150µL Nuclei Lysis Solution. The sample was inverted to mix, incubated at 56°C and 300rpm for 10 minutes, then incubated at room temperature for 20 minutes. The sample was precipitated using 150µL of Precipitation Enhancer, two DNA Capture Beads, and 700µL of isopropanol and incubated on a vertical rotating mixer at six rpm for eight minutes. The samples were then washed twice with 500µL each of Wash Buffer before incubating in 200µL 1mg/mL Proteinase K for 16 hours at room temperature. The following morning, the precipitation step was repeated with 75µL Precipitation Enhancer and 350µL isopropanol and incubated on a vertical rotating mixer at six rpm for eight minutes. The samples were washed twice more with 500µL each Wash Buffer, the beads with naked DNA were transferred to a spin column and quickly tapped on an absorbent pad to remove excess buffer, then transferred to 385µL of EB+ buffer. The DNA was allowed to solubilize at room temperature for 16 hours. The sample was poured into a bead retainer seated in a 1.5mL LoBind Eppendorf tube and spun at 16,000xg for one minute. The sample was examined and if DNA strands were visible between the flow through and the beads, the sample was spun again. The sample was mixed gently with a wide-bore pipette until maximally solubilized, pipetted roughly 10-20 times, then stored in the fridge.

10µL of sample was aspirated using a wide-bore pipette tip, with effort being made to aspirate from a homogenous region of the sample if separation is present. The 10µL was added to a 2 mL Eppendorf tube with a DNA Precipitation Bead and vortexed for two minutes to shear the DNA. The sample concentration and purity were recorded via Nanodrop.

### Library Preparation for Nanopore sequencing

Sample libraries were prepared using Oxford Nanopore’s Ultra-Long Sequencing Kit (Cat# SQK-ULK114, Batch #ULK114.20.0003). Dilute fragmentation mix was prepared using 6µL Fragmentation Mix (FRA) and 244µL Fragmentation dilution buffer (FDB) before the dilute fragmentation mix was mixed with the sample via gently pipetting with a wide-bore pipette for two minutes at room temperature. The sample was then incubated for eight more minutes at room temperature followed by 10 minutes at 75°C, five minutes on ice, and finally five minutes at room temperature. 5µL of Rapid Adaptor (RA) was added and mixed thoroughly via wide-bore pipetting and the sample was incubated at room temperature for 30 minutes. Precipitation and sample clean-up was carried out by adding a precipitation star sample followed by 500µL Precipitation Buffer (PTB). The sample was precipitated on a vertical rotating mixer for 20 minutes at 6 rpm. The sample was then spun briefly, and the supernatant was removed. The sample was eluted in 300µL Elution Buffer (EB) overnight at room temperature before being stored at 4°C.

### PromethION Loading, Reloading, and Sequencing

90µL of the sample were mixed with 100µL Sequencing Buffer (SBU) and 10µL Loading Solution Buffer (LSU) to prepare for loading onto the flow cell. This mixture was mixed thoroughly via wide-bore pipetting, incubated at room temperature for one hour, and if any insoluble DNA was visible in the sample at this point, spun down at 2,000xg for two minutes immediately before loading. The flow cell primer was prepared with 30µL Priming Tether (FTU) and 1170µL Flush Buffer (FB). The flow cell was prepared for loading by setting a P1000 pipette to 200µL, inserting the tip into the sample loading port and turning the wheel to draw up liquid until ∼10µL of liquid were visible in the pipette tip. 500µL of flow cell primer were loaded onto the flow cell. After five minutes, flow cell priming was completed by loading another 500µL of flow cell primer onto the flow cell. The sample was then immediately loaded dropwise onto the flow cell using a wide-bore pipette. After 10 minutes, sequencing was initiated. The flow cell was washed and reloaded following ONT’s wash protocol twice, as published on April 25, 2024, and reloaded following the initial loading protocol, every 24 hours.

### MinION Loading, Reloading, and Sequencing

33.8µL of the sample were mixed with 37.5µL Sequencing Buffer (SBU) and 3.7µL Loading Solution Buffer (LSU) to prepare for loading onto the flow cell. This mixture was mixed thoroughly via wide-bore pipetting, incubated at room temperature for one hour, and if any insoluble DNA was visible in the sample at this point, spun down at 2,000xg for two minutes immediately before loading. The flow cell primer was prepared with 30µL Priming Tether (FTU) and 1170µL Flush Buffer (FB). The flow cell was prepared for loading by setting a P1000 pipette to 200µL, inserting the tip into the priming port and turning the wheel to draw up liquid until ∼10µL of liquid were visible in the pipette tip. 500µL of flow cell primer was loaded onto the flow cell. After five minutes, flow cell priming was completed by loading another 500µL of flow cell primer onto the flow cell via the priming port. The sample was then immediately loaded dropwise onto the flow cell using a wide-bore pipette via the SpotON sample port. After 10 minutes, sequencing was initiated. The flow cell was washed and reloaded following ONT’s wash protocol twice, as published on April 25, 2024, and reloaded following the initial loading protocol, every 24 hours.

### Bioinformatic analysis

#### Nanopore sequencing data processing and DNAscent replication analysis

Raw Oxford Nanopore sequencing data in POD5 format were processed using a custom DNAscent analysis pipeline. First, POD5 files were converted to FAST5 using the Oxford Nanopore pod5 utility. Basecalling was performed with Guppy v6.4.2 (dna_r10.4.1_e8.2_400bps_5khz_hac configuration, GPU-enabled). Reads were aligned to the Drosophila melanogaster dm6 reference genome using Minimap2 v2.24 with the map-ont preset, followed by conversion and indexing using Samtools v1.10. DNAscent indexing was used to generate searchable indices from the FAST5 files and sequencing summary data. Replication detection was performed with DNAscent (GPU-accelerated) using a minimum read length threshold of 15 kb. ForkSense analysis was conducted with all detection modes enabled, including forks, origins, terminations, analogues, and stress signatures, using EdU and BrdU as replication markers. Bedgraph files were generated using DNAscent utility scripts to visualize replication features across the genome.

#### Track filtering and quality control for fork progression analysis

Labeled replication tracks were filtered to retain high-confidence replication fork calls. The DNAscent forkSense output BED files (leftForks_DNAscent_forkSense_stressSignatures.bed and rightForks_DNAscent_forkSense_stressSignatures.bed) were merged. Fork entries with negative DNAscent-assigned stall scores were excluded, as these indicate unreliable or ambiguous events: −1 (termination site), −2 (segmentation error), −3 (read end), and −4 (proximity to indels >100 bp). Due to the sequential EdU-then-BrdU labeling, initiation events were retained. In these cases, BrdU incorporation following EdU reliably marks fork progression from replication origins. By contrast, termination events lacking BrdU labeling likely reflect forks that completed before BrdU was incorporated and were excluded due to insufficient labeling continuity. This filtering strategy ensured that only high-confidence replication forks were included in downstream analyses.

#### BrdU and EdU track length determination, ratio calculation, and statistical analysis

Replication fork progression data were obtained from DNAscent forkSense output BED files after quality filtering. EdU and BrdU track lengths, representing base pairs synthesized during each labeling period, were extracted from these files. Entries with negative stall scores or zero EdU track length were excluded. The BrdU/EdU ratio was calculated as BrdU track length divided by EdU track length for each fork, with ratios above 100 removed as outliers. Ratio distributions were summarized by descriptive statistics and compared across groups using Mann–Whitney U tests. Ratios were analyzed alongside track lengths, stall scores, and fork direction to characterize replication dynamics.

#### Replication fork directionality (RFD) calculation

Replication fork directionality (RFD) was calculated using fork direction data from unfiltered DNAscent forkSense BED files, which contain genomic coordinates and strand annotations (“left” or “right”). The genome was divided into non-overlapping 100kb windows. For each window, the number of rightward- and leftward-moving forks was counted, and RFD was computed using the formula: RFD = (rightward − leftward) / (rightward + leftward). This yielded values ranging from −1 (fully leftward) to +1 (fully rightward), with 0 indicating balanced fork directionality.

#### Genome-wide RFD heatmap generation

Genome-wide RFD heatmaps were generated by binning RFD values into 100kb windows across all chromosomes. A matrix was constructed with chromosomes as rows and genomic bins as columns. Each cell represented the RFD value in a given genomic region. Heatmaps were visualized using a custom blue–white–red color scale centered at zero to highlight replication polarity patterns across the genome. Axes were labeled with chromosome names (y-axis) and genomic coordinates in megabases (x-axis). A color bar indicated the RFD scale.

#### Linear genomic visualization of mitochondrial replication fork tracks

DNAscent ForkSense output BED files containing BrdU-labeled mitochondrial replication tracks were filtered to include only those overlapping the mitochondrial genome. Tracks were classified by strand orientation (forward or reverse) and fork direction (left or right), then sorted by genomic start position within each category. Visualization consisted of linear plots with forward strand tracks displayed above and reverse strand tracks below a central axis, where each track is depicted as a horizontal line proportional to its genomic length. Fork direction was indicated by a color scale—blue for right-moving and orange for left-moving forks—with intensity reflecting track length.

#### Fork coverage calculation and heatmap visualization

Fork coverage was calculated by summing leftward and rightward fork counts within each non-overlapping 100kb window, based on DNAscent forkSense BED files. A fork coverage matrix was created with chromosomes as rows and genomic bins as columns, where each cell contained the total fork count for that region. Missing values were assigned to bins without any detected forks. Heatmaps were generated using the viridis color scale to represent replication fork density, with darker colors indicating higher coverage. Matrix structure and axes were standardized as described above to enable direct comparison with RFD heatmaps.

#### Origin and termination coverage heatmap generation and analysis

Replication origin or termination sites identified from DNAscent forkSense BED files were separately assigned to non-overlapping 100-kb genomic windows based on their genomic start positions. For each window, coverage was defined as the number of origins or terminations within that bin. Genome-wide coverage heatmaps were generated with chromosomes as rows and 100kb bins as columns. Origin and termination densities were visualized using continuous color scales optimized to capture dynamic range and density variation.

#### Inter-origin and inter-termination distance calculation and distribution analysis

Replication origin or termination coordinates were obtained from BED files generated by DNAscent ForkSense outputs, containing chromosome names and start and end positions. Midpoints were calculated as the average of start and end coordinates. Coordinates were grouped by chromosome and sorted by midpoint in ascending order. Inter-event distances were calculated as the difference between consecutive midpoints within the same chromosome and converted to kilobases. When plotting, outliers outside the 1st to 99th percentile range were excluded.

#### BrdU track mapping to chromatin states and comparative analysis

Previously filtered high-confidence BrdU-labeled replication tracks were used to assess replication dynamics across distinct chromatin environments. Chromatin annotations were based on a 9-state ChromHMM model, originally trained on the Drosophila dm3 genome and converted to dm6 coordinates. Regions that failed to map during the conversion process were designated as State 0. To ensure accurate chromatin state assignment, only BrdU tracks that were fully contained within a single chromatin state were included in the analysis. This was achieved using a strict containment criterion via bedtools intersect. For each retained BrdU track, we quantified: (1) the length of the BrdU-labeled region, (2) the total length of the parent DNA molecule, and (3) the ratio of BrdU length to read length as a normalized replication metric.

To simplify comparisons across different chromatin environments, the nine ChromHMM states were grouped into broader chromatin categories based on their functional characteristics. States 1–5, corresponding to transcriptionally active or accessible regions, were classified as euchromatin. States 0 and 6–8, typically associated with repressive or compact chromatin features, were categorized as heterochromatin. State 9, lacking hallmark chromatin modifications and generally considered transcriptionally inert, was labeled as a silent region. Differences between the euchromatin and heterochromatin groups were assessed using non-parametric Mann–Whitney U tests.

#### Chromatin state composition analysis of BrdU tracks

Previously chromatin state–labeled BrdU tracks were ranked by length and stratified into length-based percentile groups (Q1–Q5) to enable balanced comparisons across replication track lengths. Within each length group, overlapping BrdU tracks were merged to generate non-redundant genomic intervals, thereby avoiding redundancy bias. For each merged interval, the proportion of sequence annotated as each chromatin state was calculated. Additional normalization strategies were applied, including abundance normalization based on raw percent coverage, to facilitate relative comparisons of chromatin state biases across length groups.

BrdU-labeled replication tracks were analyzed in relation to genome-wide replication timing annotations to investigate the relationship between track length and replication timing. DNAscent ForkSense output BED files containing BrdU track coordinates were used alongside replication timing annotation files, which assign genomic intervals to four timing groups (0–3) with associated timing scores.

#### BrdU track length analysis relative to replication timing

BrdU-labeled replication tracks were analyzed in relation to genome-wide replication timing annotations to investigate the relationship between track length and replication timing. Previous filtered high-confidence ForkSense BED files containing BrdU track coordinates were used alongside replication timing annotation files, which assign genomic intervals to four timing groups (0–3) with associated timing scores. Due to the small size of replication timing segments (∼100 bp) relative to larger BrdU tracks, fixed non-overlapping 30 kb timing windows were generated across each chromosome to provide a standardized spatial framework. Each 30 kb window was assigned a dominant replication timing group based on the largest cumulative overlap with the smaller timing segments. Specifically, for each window, the total overlap length of all small timing regions belonging to each timing group was calculated, and the group with the greatest total overlap was assigned as the dominant group. This approach ensures that the timing group covering the majority of the window’s genomic space is selected. Additionally, mean timing scores and region counts—representing the average timing score and number of small timing segments overlapping the window—were computed to characterize the timing signal strength and density per window. BrdU tracks were then mapped to these timing windows by identifying overlaps. For tracks overlapping multiple windows, the window with the greatest overlap was assigned to the track, ensuring accurate spatial mapping. Track length distributions were first compared across all four replication timing groups using the Kruskal-Wallis H-test to assess overall differences. Subsequently, pairwise Mann-Whitney U tests were performed for specific group comparisons. Significance thresholds were set at p ≤ 0.05. Additionally, to explore the relationship between replication timing and replication track size, scatter plots were generated with the x-axis representing log₁₀-transformed mean replication timing scores and the y-axis representing log₁₀-transformed BrdU track lengths. This visualization highlights genome-wide trends between replication timing and track size on a continuous scale. The analysis pipeline was implemented in Python 3 using pandas for data handling, matplotlib and seaborn for visualization, and scipy for statistical testing.

#### Transcription levels for the S2 genome

PRO-seq data from Drosophila S2 cells (GSM1032758^47^), were processed to quantify gene-level transcription activity. Data containing RPKM (reads per kilobase million) values were converted from dm3 to dm6 genome builds using CrossMap BED^74^ on usegalaxy.org^75^ with dm3ToDm6.over.chain.gz from UCSC. The files were converted from bedgraph to bigWig using bedGraphToBigWig^76^. The converted data were overlapped with a BED file of all genes in the dm6 genome using UCSC’s bigWigAverageOverBed tool. This was done in a strandwise manner and genes with zero RPKM values were filtered out, resulting in a BED file containing location, strand, and RPKM values for each gene.

#### Transcriptional activity within BrdU tracks

BrdU-labeled replication tracks from filtered high-confidence forkSense BED files were analyzed alongside transcription activity (RPKM) data from Drosophila S2 cells. Tracks were divided into five length-based quantile groups (Q1–Q5). For each individual track, the mean RPKM of overlapping genes was calculated and assigned in a strand-specific manner, with each track represented as a single data point. Genes with a RPKM of 0 were excluded from this analysis. The X-axis corresponds to track length groups, and the Y-axis shows mean RPKM per track, enabling comparison of transcriptional activity across replication track lengths. Statistical differences between groups were tested using Mann–Whitney U tests. Transcription distributions were visualized with box plots and cumulative distribution functions on logarithmic scales. Analyses were conducted in Python 3 using pandas, intervaltree, scipy, matplotlib, and seaborn.

#### Analysis of replication fork length and transcriptional activity by chromatin state

Genome-wide analysis used all BrdU tracks that were not part of the chromatin state fitting and overlapped at least one gene with nonzero RPKM. For chromatin state analysis, BrdU track coordinates with chromatin state annotations were obtained from previously generated single-state fits. Only tracks overlapping at least one gene with nonzero RPKM were retained. For each track, the mean RPKM of all overlapping genes was computed. Tracks were grouped by chromatin state (0–9) and by broader functional categories: euchromatin (states 1–5), heterochromatin (states 0, 6–8), and silent intergenic (state 9). Scatter plots were generated for each group, plotting mean RPKM versus BrdU length. Linear regression lines and Pearson correlation coefficients were calculated to assess the relationship between active transcription and replication fork length.

#### Analysis of replication fork length and transcriptional activity by orientation

Replication–transcription orientation analysis was performed on BrdU-labeled replication tracks categorized as head-on or co-directional based on the direction of fork movement and overlapping gene transcription. Fork direction was obtained from DNAscent forkSense output, and gene annotations with strand and RPKM values were used to define orientation: rightward-moving forks overlapping leftward-transcribed genes (–) and leftward-moving forks overlapping rightward-transcribed genes (+) were classified as head-on, with the inverse defined as co-directional. For each track, mean RPKM of overlapping genes was calculated, and active transcription was assessed by plotting replication fork length against the mean RPKM. Scatter plots were generated separately for head-on and co-directional forks, with RPKM on the x-axis and fork length on the y-axis. Pearson correlation coefficients were computed for each group.

#### Defining origin and termination sites

Origin and termination BED files produced by DNAscent forkSense for each sequencing run were combined per run, then concatenated across runs to create comprehensive origin and termination datasets. Origin and termination events were defined by taking the midpoint of each called origin or termination event and adding 100bp to each side, resulting in a uniform set of origin and termination sites that were all 200 bps. These final center-coordinate BED files were generated for downstream analyses using Python (v3.12) and pandas (v2.2.2)^77^.

#### Replication timing data preparation

Replication timing data were retrieved from published study (GSE17280^6^). The chromosome, start, stop, and average of the three M-values for each entry were extracted and exported as a bed file. This bed file was converted from dm3 to dm6 using CrossMap^74^ BED on usegalaxy.org^75^. For conversion, the liftover file from UCSC was used (dm3ToDm6.over.chain.gz).

#### Replication timing distance analysis

Replication timing regions were grouped into quartiles, merging adjacent intervals within the same quartile to ensure coverage. The centers of adjacent inter-origin or inter-termination events are used to calculate the inter-event distance. The midpoint of each distance is then used to classify that distance into a replication timing quartile by determining the overlap of the midpoint with the replication timing bed file. Distances with midpoints that do not overlap with any replication timing region are discarded for this analysis. The distances are then grouped by quartile and the earliest and latest quartile groups are plotted using seaborn (v0.13.2) violin plots^78^. The mean and median for each quartile are calculated and the means are compared with Student’s T-test.

#### Replication timing density analysis

Origin and termination events are classified into replication timing quartiles by dividing the replication timing bed file into quartiles and adjacent reads in the same quartile are merged (this is necessary to ensure sufficient genome coverage when assigning replication quartiles based on single base overlaps). For each event, we assign the event a replication timing quartile based on the overlap of the replication timing bed file with the center of the event. Events where the center does not overlap with any region in the replication timing bed file were discarded. Once the replication timing quartile is assigned the total number of events in each quartile is summed and divided by the total genomic length mapped by that replication timing quartile.

#### Chromatin state distance analysis

The centers of adjacent events are used to calculate the inter-event distance. The midpoint of each distance is then used to classify that distance into a chromatin state by determining the overlap of the midpoint base with the chromatin state bed file. The distances in states one through five are grouped as euchromatin and the distances in states six through eight and zero are grouped as heterochromatin and plotted as violin plots. The mean and median for each group are calculated and the means are compared with Student’s T-test.

#### Chromatin state density analysis

We assign a chromatin state to each event based on the overlap of the center of that event with regions in the chromatin state bed file. If an event does not overlap with any region in the chromatin state bed file, it is discarded for this analysis. Once the chromatin state is assigned, events in states one through five are grouped as euchromatin and events in states six through eight and zero are grouped as heterochromatin events. The total number of events in each group is summed and divided by the total genomic length mapped by the chromatin state bed file for each grouping.

#### ORC2 data acquisition, preparation, and analysis

ORC2 ChIP-seq data were downloaded from GEO (GSE20887) as GFF3 files, converted to BED format, and converted from dm3 to dm6 assembly using CrossMap with UCSC chain files. The processed ORC2 peak BED file was used for downstream analyses, including inter-peak distance calculations and density comparisons substituting for origin event files. Inter-ORC2 distance is calculated as the distance between the centers of adjacent ORC2 peaks and plotted as a violin plot using seaborn.

#### Random permutation origin-ORC2 distances

For each origin, the distance to the nearest ORC2 peak is calculated and plotted as a histogram on a log_10_ scale (one is added to each distance to prevent logarithmic compute errors). A randomized imitation dataset is then generated containing the same number of events per chromosome as the true dataset but is distributed randomly along the genome. The distance from each imitated event to the nearest ORC2 peak is calculated and stored. This process of dataset imitation and distance calculation is repeated 1,000 times and the normalized results are plotted as a histogram on a log scale. The mean and median for the original and imitation dataset are calculated and a Student’s T-test was performed.

#### MNase heatmaps

MNase-seq data were obtained from GEO (GSE128689), converted from TDF to bedgraph format using igvtools tdftobedgraph (v2.17)^79^, and analyzed with deepTools^80^ computeMatrix and plotHeatmap on usegalaxy.org to visualize nucleosome occupancy centered on origins and ORC2 peaks within 2 kb windows.

#### Overlaps: expected vs. observed

Here we compute the number overlaps found within the origin dataset and compare that to the number of overlaps expected from a simulates random, independent sampling of a set of size 3,593 equal to the amount of the origin dataset (1,513). 3,593 was calculated as the expected number of mappable origins in the S2 genome given a mappable genome size of 143,726,002bps and an average inter-origin distance of 40kb. The number of repeated sampling events is recorded as the number of expected overlaps for that simulation. The simulation and sampling process are repeated 1,000 times and the mean and standard deviation of the expected number of overlaps is computed.

#### Random permutation analysis

The random permutation analysis was performed as previously described^7,60^. Briefly, ChIP-chip and ChIP-seq data sets were downloaded from modENCODE and for each factor, base pair overlap was calculated between the origin peaks or the termination peaks. For each ChIP-chip or ChIP-seq marker, we calculated the number of overlapping base-pairs (bp) between the marker and the origin/termination centers. We applied a permutation-based approach to determine whether the observed amount of overlap was more than what was expected by chance. Briefly, we calculated an empirical *p* value for the observed amount of overlap by comparing the number of overlapping bp to a simulated null distribution. We simulated the null distribution by randomly shuffling regions throughout the genome and calculating the amount of overlap for each permutation. The *p*-values were adjusted for multiple testing using the Bonferroni method.

When permuting, we matched the number and length distribution of the shuffled chromatin markers to the original set of markers and excluded all blacklisted regions from consideration [3]. For peaks obtained from ChIP-chip data, we required that the shuffled peaks maintained both the overall length distribution and the probe density of the original peak. We reshuffled any peaks were more than 0.1 units away from the original probe density until at least 99% of the original peaks were appropriately matched. We performed 1000 permutations for each marker and set of origin or termination centers.

#### UpSet diagram of Origins, ORC2, and SNS-seq

SNS-seq data was retrieved from a published study (GSE65692) and converted to dm6 using CrossMap BED on usegalaxy.org. For conversion, the liftover file from UCSC was used (dm3ToDm6.over.chain.gz). This bed file, the origins bed file, and the previously converted ORC2 ChIP-seq bed file were inputted into UpSet^81^ diagram on usegalaxy.org.

#### Nucleotide Frequency and Motif Enrichment

HOMER^82^ (v5.1) was installed locally to evaluate nucleotide bias or motif enrichment about origins. findMotifsGenome.pl was used to identify potential sequence motifs in the origin bed files. The tool was run with default inputs, with the origin bed file as the position file and dm6 as the reference genome. annotatePeaks.pl with the -annStats tag was used to determine origin enrichment in the various genomic regions. The –hist and –nuc tags were used to determine nucleotide frequency about the origins. The AC genome frequency determined using UCSC faCount (v482) tool and was plotted as dashed lines.

## DATA AVAILABILITY

All sequencing data has been deposited into GEO and will be made public upon publication.

## ACKNOWLEDGEMENTS

We thank Dave Cortez and Rahul Bhowmick for critically reading the manuscript. We are indebted to Rahul Bhowmick for sharing results prior to publication. The initial phase of this work was made possible through the National Cancer Institute (NCI) Vanderbilt-Ingram Cancer Center Support Grant (P30CA068485). This research used the Delta advanced computing and data resource which is supported by the National Science Foundation (award OAC 2005572) and the State of Illinois. Delta is a joint effort of the University of Illinois Urbana-Champaign and its National Center for Supercomputing Applications. This research was supported by NIH grant 2R35GM128650 (to J.T.N). D.S. was supported by F99AG083284 from the NIA.

## AUTHOR CONTRIBUTIONS

Conceptualization, D.H. and J.T.N; Formal analysis, D.H., C.S., M.L.B.; Investigation, D.H. and C.S.; Writing – original draft, D.H., C.S. and J.T.N.; Writing – review & editing, D.H., C.S. and J.T.N; funding acquisition, J.T.N; Supervision, J.T.N

