## Supplemental Figures for "Nanopore-based sequencing of active DNA replication reveals key principles of metazoan replication fork progression, origin and termination sites"

A

| Category | State # | Description |
|---|---|---|
| Euchromatin (Active / Transcribed) | 1 | Active Promoter / TranscriptionStart Site |
| | 2 | Actively Transcribed Exon |
| | 3 | Actively Transcribed Intron (Enhancer) |
| | 4 | Other Open Chromatin |
| | 5 | Actively Transcribed Exon on the male X chromosome (Dosage Compensated) |
| Heterochromatin | 6 | Region of Polycomb–mediated Repression |
| | 7 | Heterochromatin |
| | 8 | Heterochromatin–like region embedded in euchromatin |
| | 0 | Dm3 not-covered region (dm6–dm3) |
| Other / Unclassified | 9 | Silent Intergenic, lacking chromatin modifications |

B

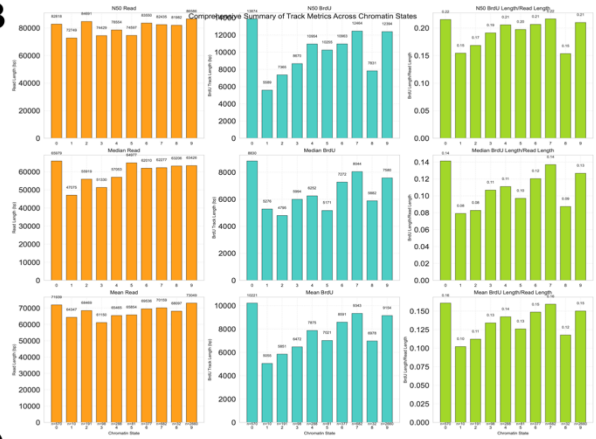


C

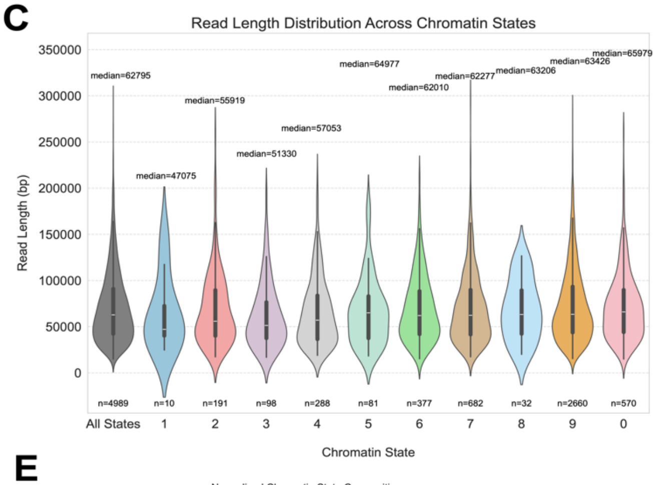


D

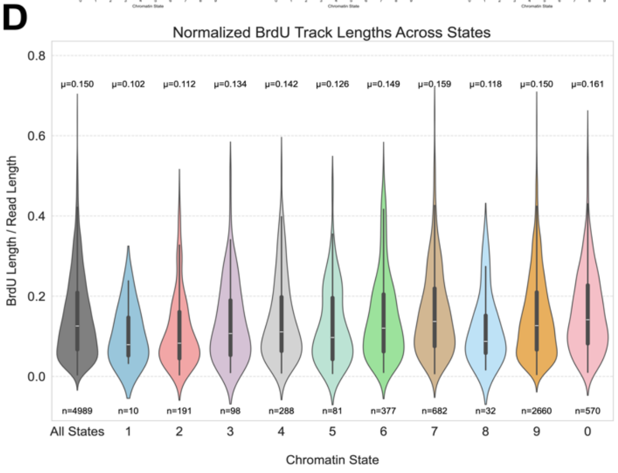


E

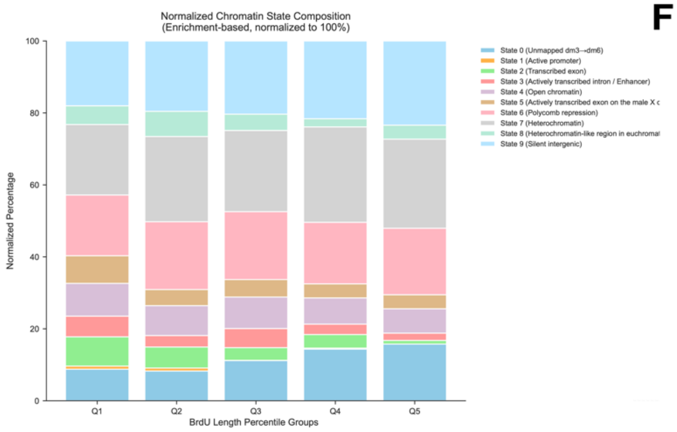


F

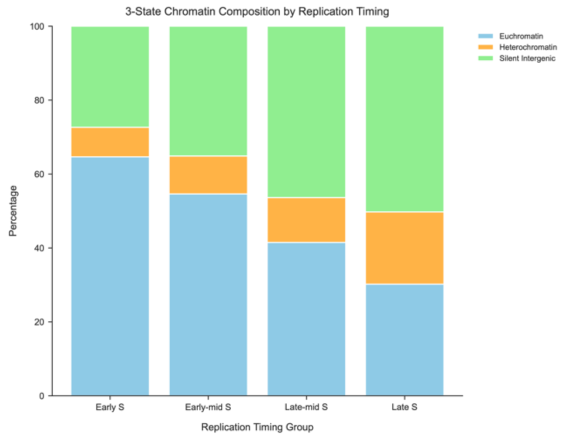


G

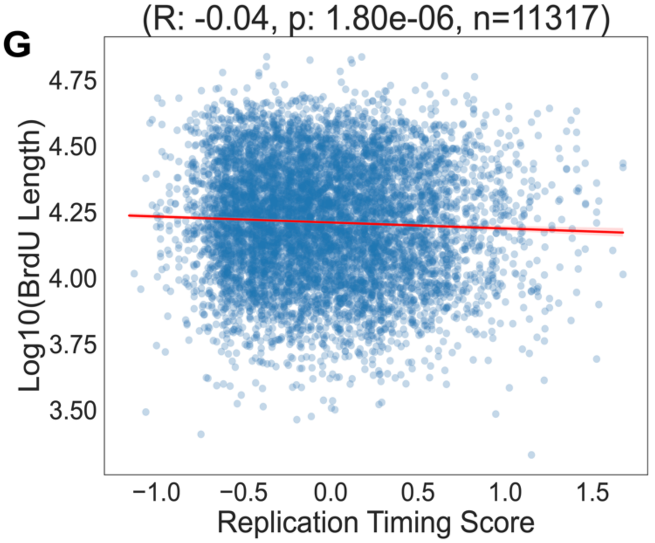

**Supplemental Figure 1. Chromatin state definitions and detailed analysis of DNAscent data**

**(A)** Legend defining the ten-state Drosophila chromatin model used in the analysis. Each state is categorized as either Euchromatin (Active / Transcribed), Heterochromatin, or Other / Unclassified, with a corresponding color and a brief description.

**(B)** Bar charts summarizing various metrics for BrdU track analysis across the ten chromatin states. The panels show the average read length, average BrdU track length, and the ratio of BrdU track length to read length.

**(C)** Violin plots showing the distribution of total read lengths (in base pairs, bps) for reads that map to the different chromatin states. The first violin represents all reads combined ("All States"). The median read length for each state is shown above the corresponding violin.

**(D)** Violin plots showing the distribution of normalized BrdU track lengths, calculated as the ratio of BrdU track length to total read length for each read. The median for each chromatin state is shown above the corresponding violin.

**(E)** Stacked bar chart showing the chromatin state composition within different BrdU length percentile groups (Q1-Q5).

**(F)** Stacked bar chart showing the composition of a simplified 3-state chromatin model (Euchromatin, Heterochromatin, Silent Intergenic) across different replication timing groups (Early S, Early-mid S, Late-mid S, Late S).

**(G)** Scatter plot showing the correlation between $\log_{10}$-transformed BrdU track lengths and the replication timing score. Each blue dot represents a single BrdU-labeled read. The red line indicates the best-fit linear regression (Pearson's $R = -0.04$; $p = 1.8 \times 10^{-6}$; $n = 11{,}317$).

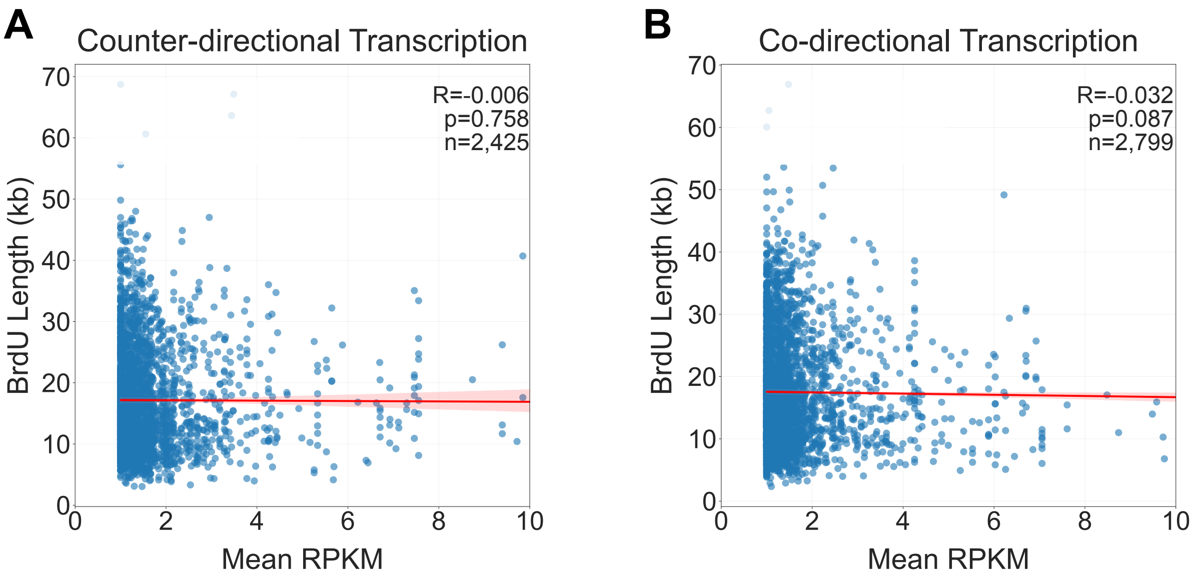


**Supplemental Figure 2. Correlation between BrdU length and active transcription levels based on relative orientation.**

**(A)** Scatter plot showing the correlation between BrdU track length and RPKM values for replication forks moving counter-directionally (head-on) to transcription. The red line indicates the linear regression fit (Pearson's r = 0.006; p = 7.58 x $10^{-1}$; N = 2,425).

**(B)** Scatter plot showing the correlation between BrdU track length and RPKM values for replication forks moving co-directionally with transcription. The red line indicates the linear regression fit (Pearson's r = -0.032; p = 8.7 x $10^{-2}$; N = 2,799).

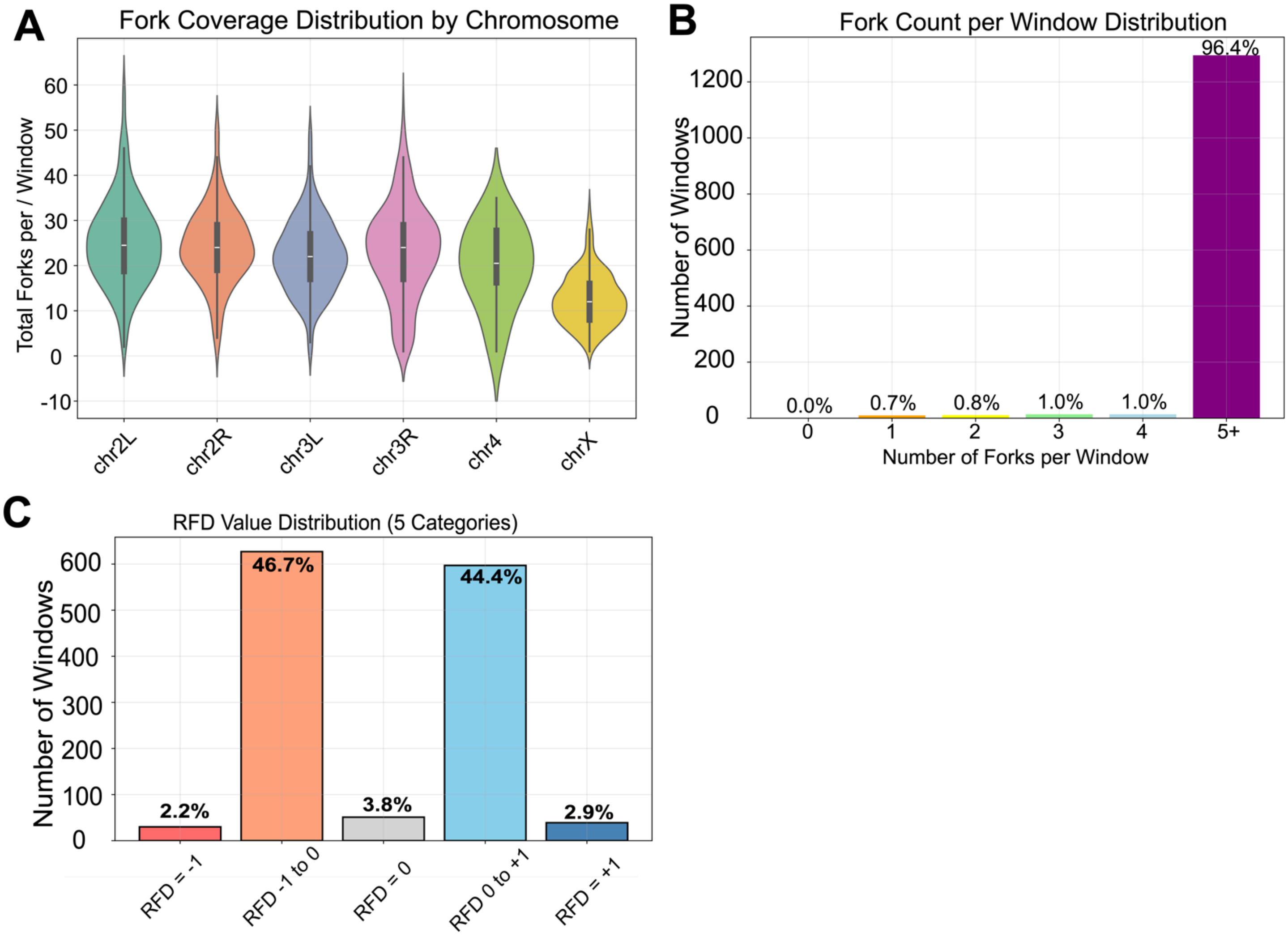


**Supplemental Figure 3. Analysis of replication fork distribution and directionality.**

**(A)** Violin plot showing the distribution of the total number of replication forks per 100kb window across different Drosophila chromosomes. The line within each violin indicates the median.

**(B)** Bar chart showing the distribution of windows based on the number of replication forks they contain. The percentages of windows for each fork count are indicated above the bars.

**(C)** Bar chart showing the distribution of windows across five categories of Replication Fork Directionality (RFD) values. The percentage of windows within each RFD category is indicated above the bars.

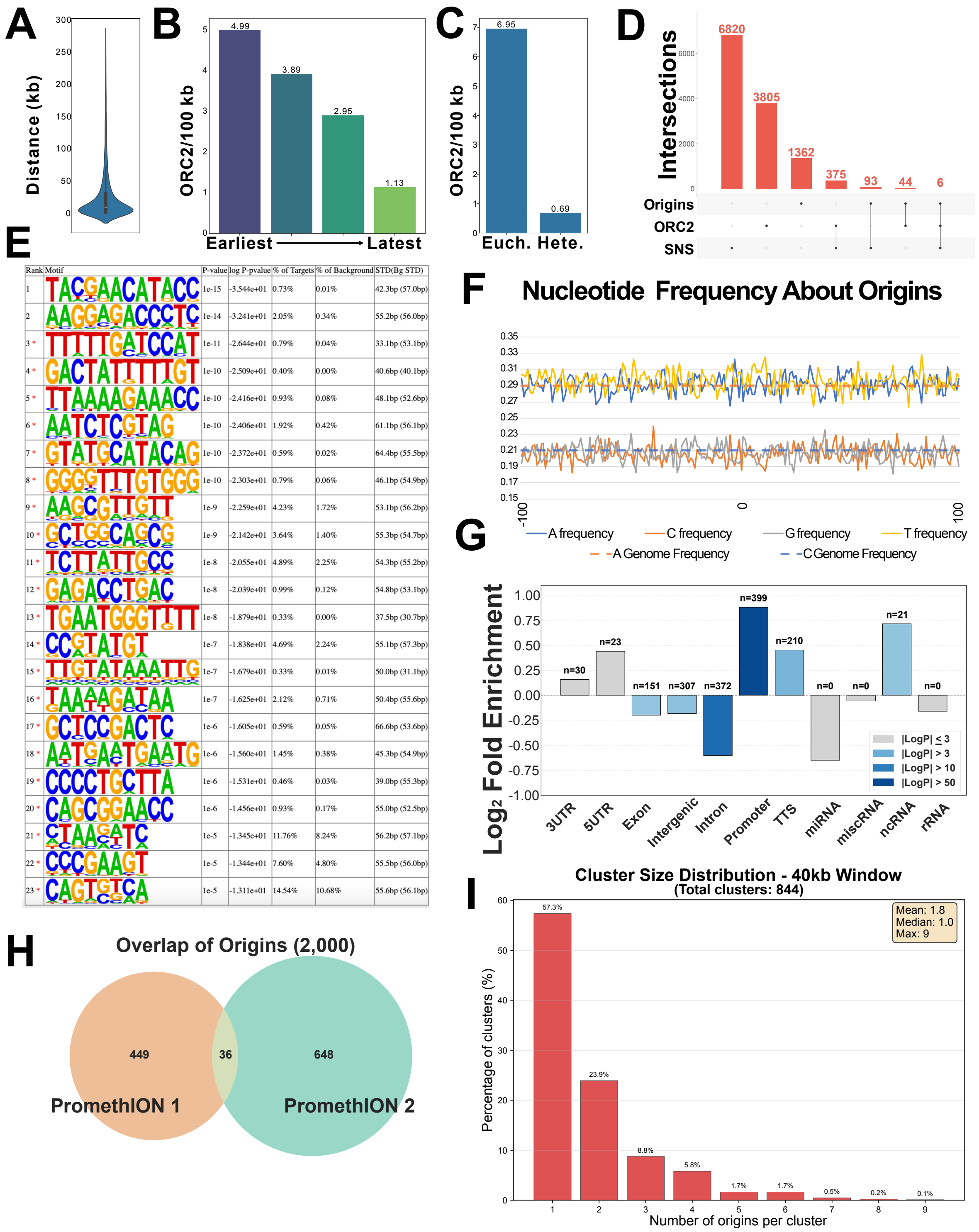


| Rank | Motif | P-value | log P-pvalue | % of Targets | % of Background | STD(Bg STD) |
|---|---|---|---|---|---|---|
| 1 | TACGAACATACC | 1e-15 | -3.544e+01 | 0.73% | 0.01% | 42.3bp (57.0bp) |
| 2 | AAGGAGACCCTC | 1e-14 | -3.241e+01 | 2.05% | 0.34% | 55.2bp (56.0bp) |
| 3 * | TTTTTGATCCAT | 1e-11 | -2.644e+01 | 0.79% | 0.04% | 33.1bp (53.1bp) |
| 4 * | GACTATTTTTGT | 1e-10 | -2.509e+01 | 0.40% | 0.00% | 40.6bp (40.1bp) |
| 5 * | TTAAAAGAAACC | 1e-10 | -2.416e+01 | 0.93% | 0.08% | 48.1bp (52.6bp) |
| 6 * | AATCTCGTAG | 1e-10 | -2.406e+01 | 1.92% | 0.42% | 61.1bp (56.1bp) |
| 7 * | GTATGCATACAG | 1e-10 | -2.372e+01 | 0.59% | 0.02% | 64.4bp (55.5bp) |
| 8 * | GGGGTTTGTGGG | 1e-10 | -2.303e+01 | 0.79% | 0.06% | 46.1bp (54.9bp) |
| 9 * | AAGCGTTGTT | 1e-9 | -2.259e+01 | 4.23% | 1.72% | 53.1bp (56.2bp) |
| 10 * | GCTGGCAGCG | 1e-9 | -2.142e+01 | 3.64% | 1.40% | 55.3bp (54.7bp) |
| 11 * | TCTTATTGCC | 1e-8 | -2.055e+01 | 4.89% | 2.25% | 54.3bp (55.2bp) |
| 12 * | GAGACCTGAC | 1e-8 | -2.039e+01 | 0.99% | 0.12% | 54.8bp (53.1bp) |
| 13 * | TGAATGGGTTTT | 1e-8 | -1.879e+01 | 0.33% | 0.00% | 37.5bp (30.7bp) |
| 14 * | CCGTATGT | 1e-7 | -1.838e+01 | 4.69% | 2.24% | 55.1bp (57.3bp) |
| 15 * | TTGTATAAATTG | 1e-7 | -1.679e+01 | 0.33% | 0.01% | 50.0bp (31.1bp) |
| 16 * | TAAAAGATAA | 1e-7 | -1.625e+01 | 2.12% | 0.71% | 50.4bp (55.6bp) |
| 17 * | GCTCCGACTC | 1e-6 | -1.605e+01 | 0.59% | 0.05% | 66.6bp (53.6bp) |
| 18 * | AATGAATGAATG | 1e-6 | -1.560e+01 | 1.45% | 0.38% | 45.3bp (54.9bp) |
| 19 * | CCCCTGCTTA | 1e-6 | -1.531e+01 | 0.46% | 0.03% | 39.0bp (55.3bp) |
| 20 * | CAGCGGAACC | 1e-6 | -1.456e+01 | 0.93% | 0.17% | 55.0bp (52.5bp) |
| 21 * | CTAAGATC | 1e-5 | -1.345e+01 | 11.76% | 8.24% | 56.2bp (57.1bp) |
| 22 * | CCCGAAGT | 1e-5 | -1.344e+01 | 7.60% | 4.80% | 55.5bp (56.0bp) |
| 23 * | CAGTGTCA | 1e-5 | -1.311e+01 | 14.54% | 10.68% | 55.6bp (56.1bp) |

**Supplemental Figure 4. ORC2 Peak distribution and chromatin context analyses.**

**(A)** Violin plot showing the distribution of inter-ORC2 distances (in kilobases, kb) for all ORC2 binding peaks in the dataset (n=4,230), with a median distance of 9.85 kb.

**(B)** Bar chart showing ORC2 binding site density (peaks per 100 kb) between early and late replicating domains.

**(C)** Bar chart showing ORC2 binding site density (peaks per 100 kb) between euchromatin and heterochromatin.

**(D)** UpSet diagram showing the number of overlaps between identified replication origins, ORC2 binding sites, and origins identified by SNS-seq.

**(E)** Line plot showing the average nucleotide frequency around origins as solid lines (adenine: blue, cytosine: orange, guanine: grey, thymidine: yellow). The average nucleotide frequency is shown as dashed lines (adenine: dashed orange, cytosine: dashed blue).

**(F)** HOMER Motif analysis showing no sequence motif enrichment within origins. Red stars indicate a likely false positive.

**(G)** HOMER Annotate Peaks analysis showing the $log_2$-fold enrichment of origins within genomic feature. The plot shows that origins are enriched at promoters and transcription termination sites (TTS) and depleted in gene bodies (exons and introns).

**(H)** Venn diagram of the overlap of identified origins at ten-times the window sized used for analysis (size = 2,000bps) between two biological replicates, "PromethION 1" and "PromethION 2".

**(I)** Distribution of origin sites within 40kb windows containing one or more origins

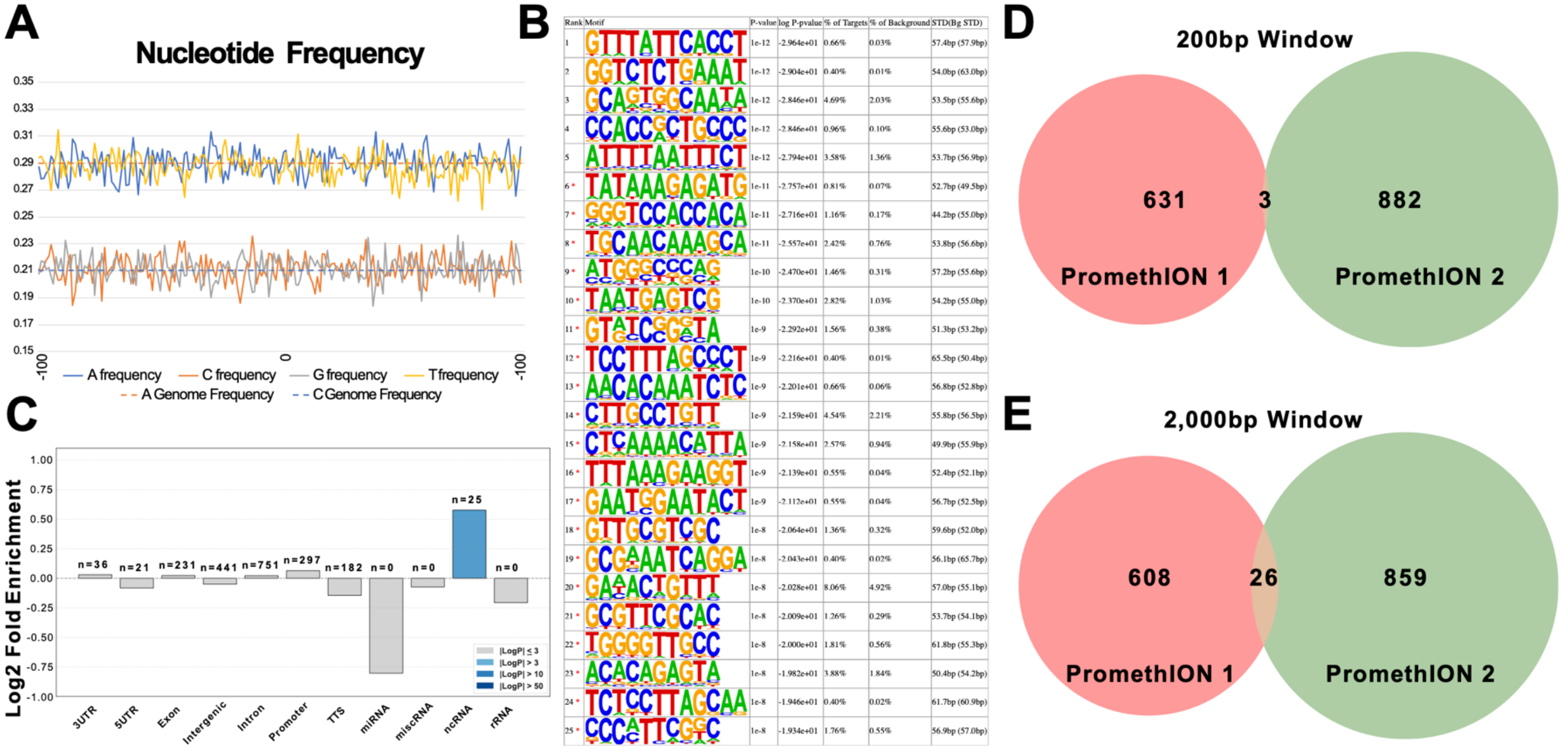


**Supplemental Figure 5: Termination Enrichment and Overlap**

**(A)** Line plot showing the average nucleotide frequency around termination events as solid lines(adenine: blue, cytosine: orange, guanine: grey, thymidine: yellow). The average nucleotide frequency is shown as dashed lines (adenine: dashed orange, cytosine: dashed blue).

**(B)** HOMER Motif analysis showing no sequence motif enrichment within terminations. Red stars indicate a likely false positive.

**(C)** HOMER Annotate Peaks analysis showing the $log_2$-fold enrichment of terminations within various genomic features relative to the genome background. The plot shows significant enrichment at noncoding RNA sites.

**(D)**Venn diagram illustrating the overlap of identified terminations at window sizes 200bps between two biological replicates.

**(E)** Venn diagram illustrating the overlap of identified terminations at window sizes 2000bps between two biological replicates.
